# Epithelial sensing of vitamin A shapes intestinal antimicrobial defense

**DOI:** 10.64898/2026.03.08.710399

**Authors:** Gabriella Quinn, Daniel C. Propheter, Kartik Kulkarni, Matthew Johnson, Gonçalo Vale, Jeffrey G. McDonald, Ann Johnson, Brian Hassell, Cassie L. Behrendt, Nikhil V. Munshi, Lora V. Hooper

## Abstract

Vitamin A is a central regulator of intestinal adaptive immunity, but its role in innate immunity is less defined. Antimicrobial proteins form a chemical barrier that protects the intestinal epithelium from microbial invasion. Among these, REG3 family lectins are induced by the microbiota, yet how nutritional cues intersect with microbial signals to control their expression remains unclear. Here, we show that dietary vitamin A promotes expression of REG3 antimicrobial lectins, including REG3G, in intestinal epithelial cells from both mice and humans. This induction is mediated by retinoic acid and requires retinoic acid receptor (RAR) signaling. Mechanistically, RARs bind directly to the *Reg3g* promoter adjacent to a STAT3 binding site. As STAT3 mediates microbiota-induced IL-22 signaling in epithelial cells, this arrangement provides a molecular framework for integrating nutritional and microbial inputs at the level of REG3G transcription. Extending these findings, we demonstrate that vitamin A–retinoic acid signaling similarly promotes expression of α-defensin antimicrobial proteins. Together, these findings define a transcriptional mechanism by which vitamin A enhances epithelial antimicrobial defenses and strengthens mucosal innate immunity.

**Graphical Abstract:** 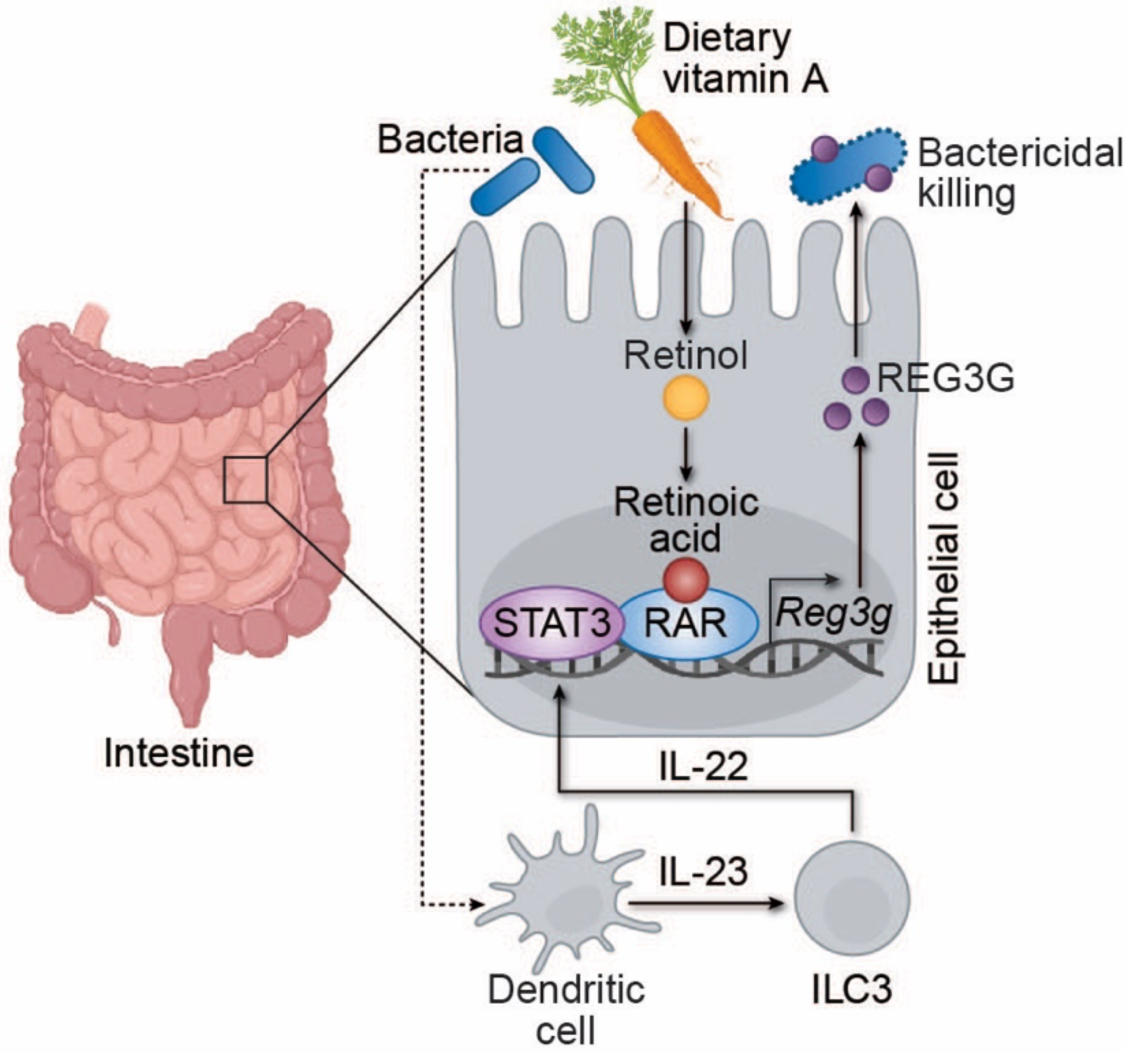

**Highlights:**

- Vitamin A promotes epithelial expression of REG3 antimicrobial proteins in the intestine
- Retinoic acid receptors (RARs) directly activate mouse *Reg3g* and human *REG3G* transcription
- RARs bind the *Reg3g* promoter adjacent to STAT3, integrating nutritional and microbial signals
- Vitamin A–RAR signaling broadly regulates epithelial antimicrobial programs, including α-defensins

## Introduction

The intestine is continuously exposed to dense microbial communities, requiring robust barrier defenses that preserve host-microbe mutualism. This protection depends on a coordinated mucosal immune system composed of immune cells and the intestinal epithelium. Beyond serving as a physical barrier, the epithelium actively defends the host by secreting antimicrobial proteins that create a chemical shield between luminal microbes and host tissues. These proteins constitute a first line of mucosal defense by disrupting bacterial membranes.^1,2^ Among the best characterized are REG3 lectins, which preferentially target Gram-positive bacteria, and α-defensins, which act primarily against Gram-negative species.^3^

REG3 lectins are distinctive in that their expression is induced by microbial cues.^4^ This pathway is well-defined: microbe-associated molecular patterns activate dendritic cells in the lamina propria, which stimulate group 3 innate lymphoid cells to produce interleukin-22 (IL-22).^5,6^ IL-22 in turn activates the epithelial transcription factor STAT3, driving *Reg3b* and *Reg3g* expression.^7–9^ In contrast, how nutritional signals influence epithelial antimicrobial programs remains largely unknown.

Vitamin A is a lipid-soluble micronutrient that plays a central role in adaptive immune responses to mucosal infection.^10–17^ Dendritic cells metabolize vitamin A into retinoic acid (RA), which shapes T– and B–cell differentiation over the course of days.^13,18–21^ This delayed, immune cell–centered mechanism raises the possibility that vitamin A may also exert more immediate, local effects at the epithelial barrier, potentially by regulating antimicrobial protein expression. Whether vitamin A directly controls antimicrobial programs in intestinal epithelial cells, however, is unclear.

Here, we show that vitamin A promotes antimicrobial gene expression in intestinal epithelial cells in both mice and human model systems. Transcripts encoding REG3 family members are among the most strongly induced, revealing a link between vitamin A and epithelial innate immunity. Using complementary in vitro and in vivo approaches, we demonstrate that this response is mediated by RA and requires retinoic acid receptor (RAR) signaling. RARs bind to the *Reg3g* promoter adjacent to a STAT3 binding site, providing a molecular framework for integration of nutritional and microbial signals at the level of transcription. Extending this mechanism, we show that vitamin A–RA signaling promotes expression of α-defensin antimicrobial proteins. Together, these findings define a mechanism by which vitamin A regulates epithelial antimicrobial programs and promotes first-line mucosal innate defense.

## Results

### Vitamin A enhances epithelial expression of REG3 antimicrobial proteins in the small intestine

Vitamin A is a key regulator of intestinal adaptive immunity^10,11,13^, but its role in epithelial innate immunity is less defined. To examine this role, we used a mouse model of dietary vitamin A deprivation in which dams were fed a vitamin A–deficient diet during gestation and lactation, and offspring were maintained on the deficient diet into adulthood (Fig. 1A).^12,22,23^ As expected, these mice exhibited markedly reduced serum retinol levels, confirming systemic deficiency (Fig. S1A).

**Figure 1.**
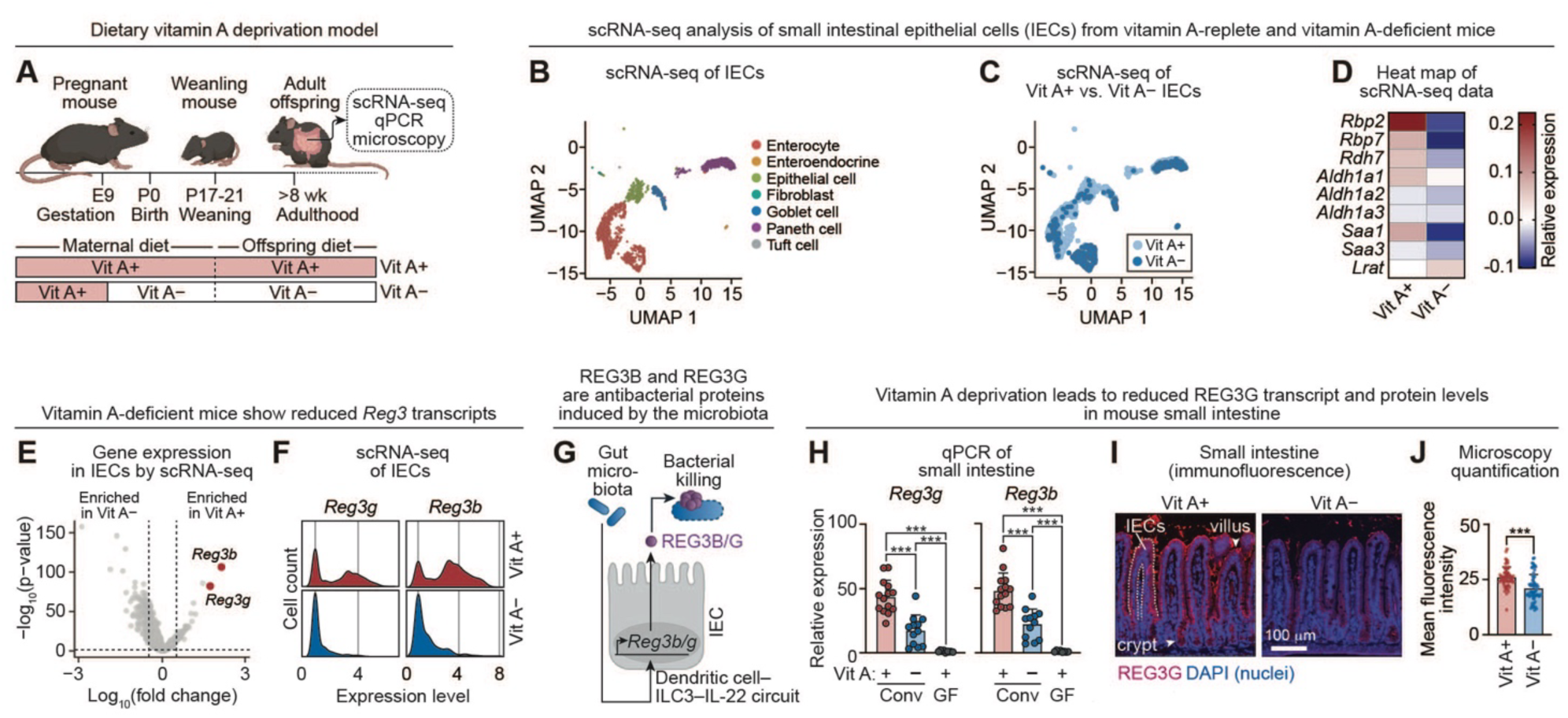
Vitamin A enhances epithelial expression of REG3 antimicrobial proteins in the small intestine. **(A)** Vitamin A deprivation model. Pregnant dams were fed vitamin A–deficient (Vit A–) or vitamin A–replete (Vit A+) diets during gestation. Offspring were maintained on the maternal diet for at least 8 weeks. **(B)** Single cell RNA sequencing (scRNA-seq) was performed on a single cell suspension from the mouse small intestine. Live cells were bar-coded and sequenced, and cells annotated as epithelial cells were analyzed and are displayed as a two-dimensional reduction plot (UMAP). Both Vit A+ and Vit A– cells are plotted (n=3 mice per group). **(C)** Distribution of epithelial cell subsets from Vit A+ and Vit A– mice (n=3) projected on the same UMAP as in (B). **(D)** Heatmap of average normalized expression of known vitamin A-responsive genes from the scRNA-seq analysis of IECs. **(E)** Volcano plot of vitamin A-dependent changes in gene expression in IECs analyzed by scRNA-seq. *Reg3b* and *Reg3g* are highlighted in red. **(F)** Density plots of *Reg3b* and *Reg3g* expression in IECs analyzed by scRNA-seq. **(G)** The gut microbiota induces expression of the antibacterial proteins REG3β and REG3γ. Microbial molecular patterns activate a dendritic cell–ILC3 signaling relay that drives IL-22 production, which induces *Reg3b* and *Reg3g* expression in IECs. **(H)** qPCR analysis of *Reg3g* and *Reg3b* transcript abundance in small intestines from conventional mice fed Vit A+ (n=12) and Vit A– (n=14) diets, and germ-free mice fed a Vit A+ diet (n=9). Mice were from three independent litters. Each data point represents one mouse. **(I)** Immunofluorescence microscopy of REG3G in the distal small intestine of Vit A+ or Vit A– mice. Sections were stained for REG3G (red) and counterstained with DAPI (blue). Scale bar, 100 μm. Images are representative of three mice per group. **(J)** Mean fluorescence intensities of at least 65 villi were determined across three mice in each dietary group. Vit A+, vitamin A+; Vit A–, vitamin A–; wk, weeks; scRNA-seq, single cell RNA sequencing; IEC, intestinal epithelial cell; UMAP, Uniform Manifold Approximation and Projection; qPCR, quantitative real-time PCR; REG3B, regenerating islet-derived protein 3β; REG3G, regenerating islet-derived protein 3γ; Conv, conventional; GF, germ-free. Means ± SEM are plotted; ***p < 0.001 by Mann-Whitney test. See also Figure S1.

To assess how vitamin A influences epithelial gene expression, we performed single-cell RNA sequencing on small intestinal cell suspensions enriched for live intestinal epithelial cells (IECs) (Fig. 1B). Cell identities were assigned by mapping to the Mouse Cell Atlas of the adult small intestine (Fig. S1B).^24^ Vitamin A–replete and vitamin A–deficient mice displayed similar epithelial cell-type distributions, with the most pronounced transcriptional differences observed in enterocytes (Fig. 1C). As expected, epithelial cells from vitamin A–deficient mice showed reduced expression of genes involved in vitamin A metabolism and transport (Fig. 1D). Differential expression analysis identified *Reg3b* and *Reg3g* among the most significantly downregulated transcripts (Fig. 1E). Single-cell analysis further revealed both a reduced fraction of *Reg3*-expressing cells and decreased transcript abundance per cell in vitamin A–deficient mice (Fig. 1F). Because the microbiota induces *Reg3b/g* transcription through a dendritic cell–ILC3 –IL-22–STAT3 circuit (Fig. 1G),^4–6,9^ these findings suggest that vitamin A amplifies microbiota-driven antimicrobial gene expression.

We validated these observations by qPCR of small intestinal tissue from three independent litters, confirming reduced *Reg3b* and *Reg3g* transcript levels in vitamin A–deficient mice (Fig. 1H). This decrease was accompanied by diminished REG3B and REG3G protein abundance (Fig. 1I, J; Fig. S1C). The reduction in REG3 expression was not attributable to altered numbers of Paneth cells, which secrete abundant antimicrobial proteins (Fig. S1D, E). Although *Reg3* transcript levels were decreased in vitamin A–deficient mice, they remained higher than in germ-free animals, indicating that both microbial signals and vitamin A are required for maximal *Reg3* induction.

REG3G and other antimicrobial proteins maintain spatial segregation between the epithelium and luminal bacteria.^25,26^ We therefore hypothesized that vitamin A deficiency might permit increased microbial proximity to the intestinal epithelial surface. To test this prediction, we gavaged mice with a microbiota capable of stimulating epithelial antimicrobial responses.^27^ Bacterial localization was assessed three days later, a time frame that predominantly reflects innate responses. Consistent with diminished antimicrobial activity, vitamin A–deficient mice exhibited increased microbial encroachment at the epithelial surface (Fig. S1F). Together, these findings indicate that dietary vitamin A reinforces microbiota-induced REG3 expression and promotes epithelial barrier integrity in the small intestine.

### The vitamin A metabolite retinoic acid promotes epithelial REG3G expression

We next investigated how vitamin A promotes REG3G expression. Because *Reg3g* and *Reg3b* are coordinately regulated in mice^5^, we focused on REG3G as a representative family member. To determine whether vitamin A–dependent induction occurs directly in epithelial cells, we established an in vitro model using HT-29 human intestinal epithelial cells. Cells were cultured in charcoal-stripped media to deplete vitamin A and related lipophilic compounds.^28,29^ Depletion was confirmed by measurement of cellular retinol, the vitamin A metabolite that is the precursor to RA (Fig. S2). Cells were then treated with IL-22 in the presence or absence of exogenous retinol. Consistent with our in vivo findings, *REG3G* expression was minimal without IL-22, increased upon IL-22 stimulation, and was further augmented by retinol (Fig. 2A). These results indicate that vitamin A enhances *REG3G* expression through an epithelial cell–intrinsic mechanism conserved between mice and humans.

**Figure 2.**
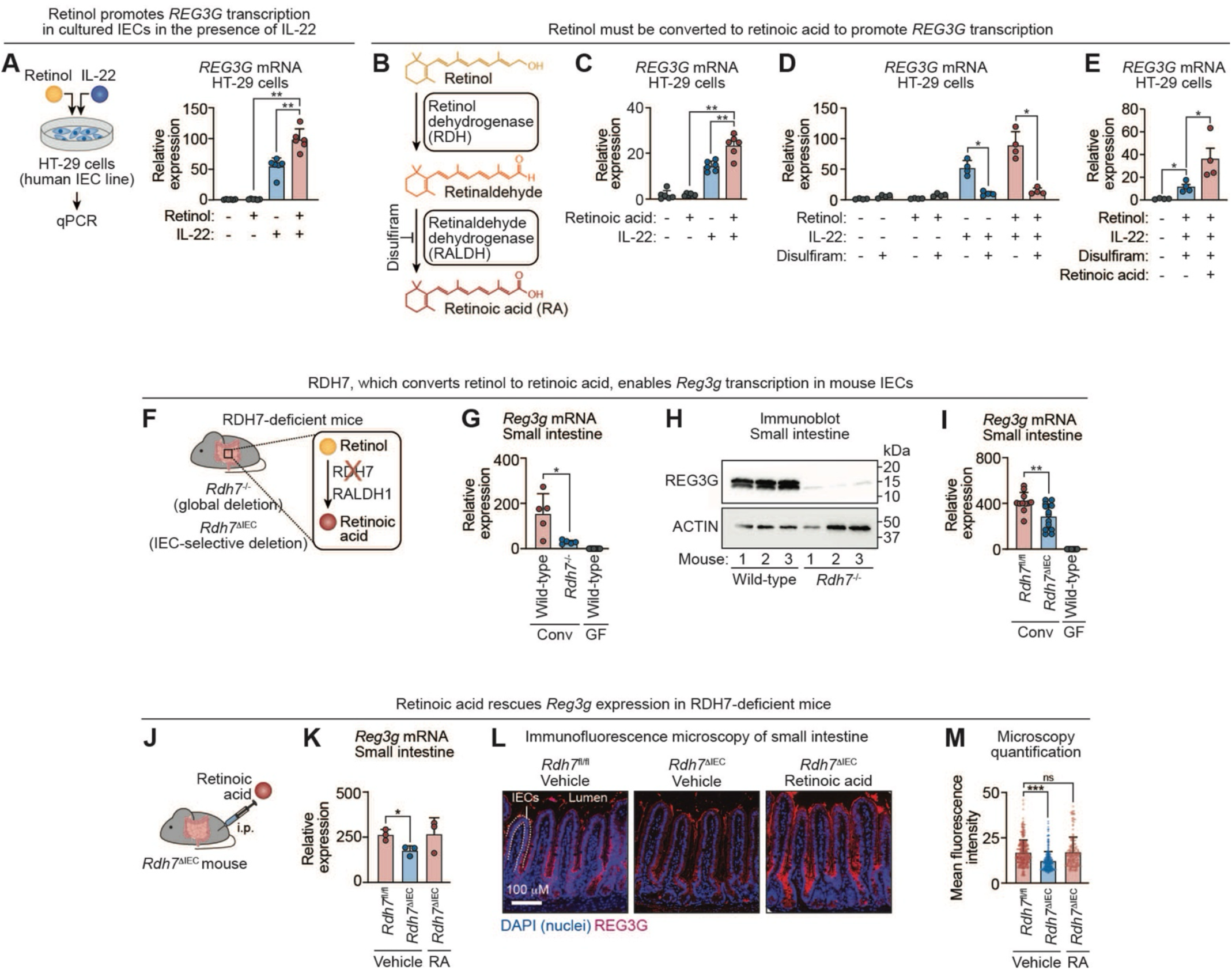
The vitamin A metabolite retinoic acid promotes epithelial REG3G expression. **(A)** HT-29 cells were treated with 1 *μ*M retinol and/or 100 ng/mL IL-22. *REG3G* transcripts were quantified by qPCR 18 hours later. Each data point represents one experimental replicate (n=6 per group). **(B)** Retinol is converted to RA through a two-step enzymatic reaction catalyzed by retinol/alcohol dehydrogenases and retinaldehyde/aldehyde dehydrogenases. Disulfiram inhibits aldehyde dehydrogenase enzymatic activity (including RALDH). **(C)** HT-29 cells were treated with 1 *μ*M RA and/or 100 ng/mL IL-22 in the presence of vehicle or 100 *μ*M disulfiram. *REG3G* transcripts were quantified by qPCR after 18 hours. Each data point represents an independent experimental replicate (n=6 per group). **(D)** qPCR analysis of *REG3G* transcripts in HT-29 cells treated with retinol and IL-22 in the presence or absence of disulfiram. Each data point represents one experimental replicate (n=4 per group). **(E)** qPCR analysis of *REG3G* transcripts in HT-29 cells treated with retinol and IL-22 in the presence of disulfiram, with rescue by RA. Each data point represents an independent experimental replicate (n=4 per group). **(F)** The *Rdh7*^-/-^ mouse carries a global deletion of the gene encoding RDH7, which catalyzes the first step in the retinol-to-RA conversion. *Rdh7*^μIEC^ mice harbor an IEC–specific deletion of *Rdh7*, generated by crossing *Rdh7*^fl/fl^ mice with Villin-Cre transgenic mice. **(G)** qPCR analysis of *Reg3g* transcripts in the small intestines of conventional wild-type (n=5) and *Rdh7*^-/-^ (n=5) mice, and germ-free wild-type (n=21) mice. Each data point represents one mouse. **(H)** Immunoblot of REG3G in small intestines from conventional wild-type (n=3) and *Rdh7*^-/-^ (n=3) mice. ACTIN was the loading control. Each lane is from one mouse. **(I)** qPCR analysis of *Reg3g* transcripts in small intestines from six litters of conventional *Rdh7*^fl/fl^ (n=11) and *Rdh7*^μIEC^ (n=13) mice and germ-free wild-type (n=21) mice. Each data point represents one mouse. **(J)** *Rdh7*^μIEC^ mice received two intraperitoneal injections of vehicle or retinoic acid (1 *μ*M), administered 12 hours apart. Mice were sacrificed 12 hours after the last injection. **(K)** qPCR analysis of *Reg3g* transcripts in the intestines of two litters of *Rdh7*^fl/fl^ (n=3) and *Rdh7*^μIEC^ (n=6) littermates injected intraperitoneally with RA or vehicle. Each data point represents one mouse. **(L)** Immunofluorescence microscopy of REG3G in the small intestines of *Rdh7*^fl/fl^ and *Rdh7*^μIEC^ littermates injected via the intraperitoneal route with retinoic acid (RA) or vehicle. Sections were stained for REG3G (red) and counterstained with DAPI (blue). Scale bar, 100 μm. Images are representative of at least three fields per sample from two independent experiments (three littermates per group). **(M)** Mean fluorescence intensities of at least 150 villi from the images represented in (L) were quantified across at least two mice in each experimental group. IEC, intestinal epithelial cell; REG3G, regenerating islet-derived protein 3γ; qPCR, quantitative real-time PCR; RDH, retinol dehydrogenase; RALDH, retinaldehyde dehydrogenase; Conv, conventional; GF, germ-free; i.p., intraperitoneal; RA, retinoic acid. Means ± SEM are plotted; *p < 0.05; **p < 0.01; ***p < 0.001; ns, not significant by Mann-Whitney test. See also Figure S2.

Retinol is not itself a transcriptional ligand. Instead, vitamin A–dependent transcription is mediated by RA, which is generated intracellularly from retinol through sequential reactions catalyzed by retinol dehydrogenases (RDHs) and retinaldehyde dehydrogenases (RALDHs) (Fig. 2B). To determine whether RA is sufficient to promote *REG3G* transcription, we added RA directly to HT-29 cells in the presence of IL-22. Like retinol, RA enhanced IL-22–dependent *REG3G* expression (Fig. 2C). In contrast, pharmacologic inhibition of aldehyde dehydrogenase activity with disulfiram reduced retinol-dependent *REG3G* induction, and this effect was rescued by exogenous RA (Fig. 2D, E). Thus, conversion of retinol to RA is required for vitamin A–dependent enhancement of *REG3G* transcription.

Disulfiram also attenuated IL-22–induced *REG3G* expression in the absence of added retinol (Fig. 2D). This effect may reflect residual but undetectable intracellular vitamin A stores despite culture in charcoal-stripped media. Alternatively, because disulfiram broadly inhibits aldehyde dehydrogenases, these enzymes may contribute to IL-22–dependent *REG3G* induction independent of RA synthesis.

We next tested whether epithelial conversion of retinol to RA is required for REG3G expression in vivo. Retinol dehydrogenase 7 (RDH7) is the predominant retinol dehydrogenase expressed by intestinal epithelial cells (Fig. 2F).^30^ Global deletion of *Rdh7* resulted in reduced *Reg3g* transcripts and decreased REG3G protein abundance in the small intestine compared with wild-type controls (Figure 2F–H). A similar reduction was observed in mice with intestinal epithelial cell-specific deletion of *Rdh7* (*Rdh7*^μIEC^), indicating that epithelial RDH7 supports *Reg3g* expression (Fig. 2I). Importantly, intraperitoneal administration of RA restored *Reg3g* transcript levels and rescued epithelial REG3G protein expression in *Rdh7*^μIEC^ mice (Fig. 2J–M). Together, these findings demonstrate that RA generated within intestinal epithelial cells is required to promote REG3G expression in vivo.

### Retinoic acid receptors drive transcription of mouse *Reg3g* and human *REG3G* genes

RA regulates gene transcription through RARs, ligand-activated transcription factors that bind retinoic acid response elements (RAREs) in target gene promoters (Fig. 3A).^31–33^ The mouse genome encodes three RAR paralogs: RARα (RARA), RARβ (RARB), and RARγ (RARG). To determine whether RAR activity is required for RA-dependent *REG3G* expression, we treated HT-29 human intestinal epithelial cells with the pan-RAR antagonist BMS493. In the presence of IL-22 and RA, BMS493 markedly reduced *REG3G* transcript abundance (Fig. 3B). Conversely, the pan-RAR agonist Ch55 increased *REG3G* transcription even in the absence of RA (Fig. 3B), indicating that RAR activation is sufficient to drive *REG3G* expression. These pharmacologic data suggest that RAR activity is required for RA-dependent *REG3G* transcription.

**Figure 3.**
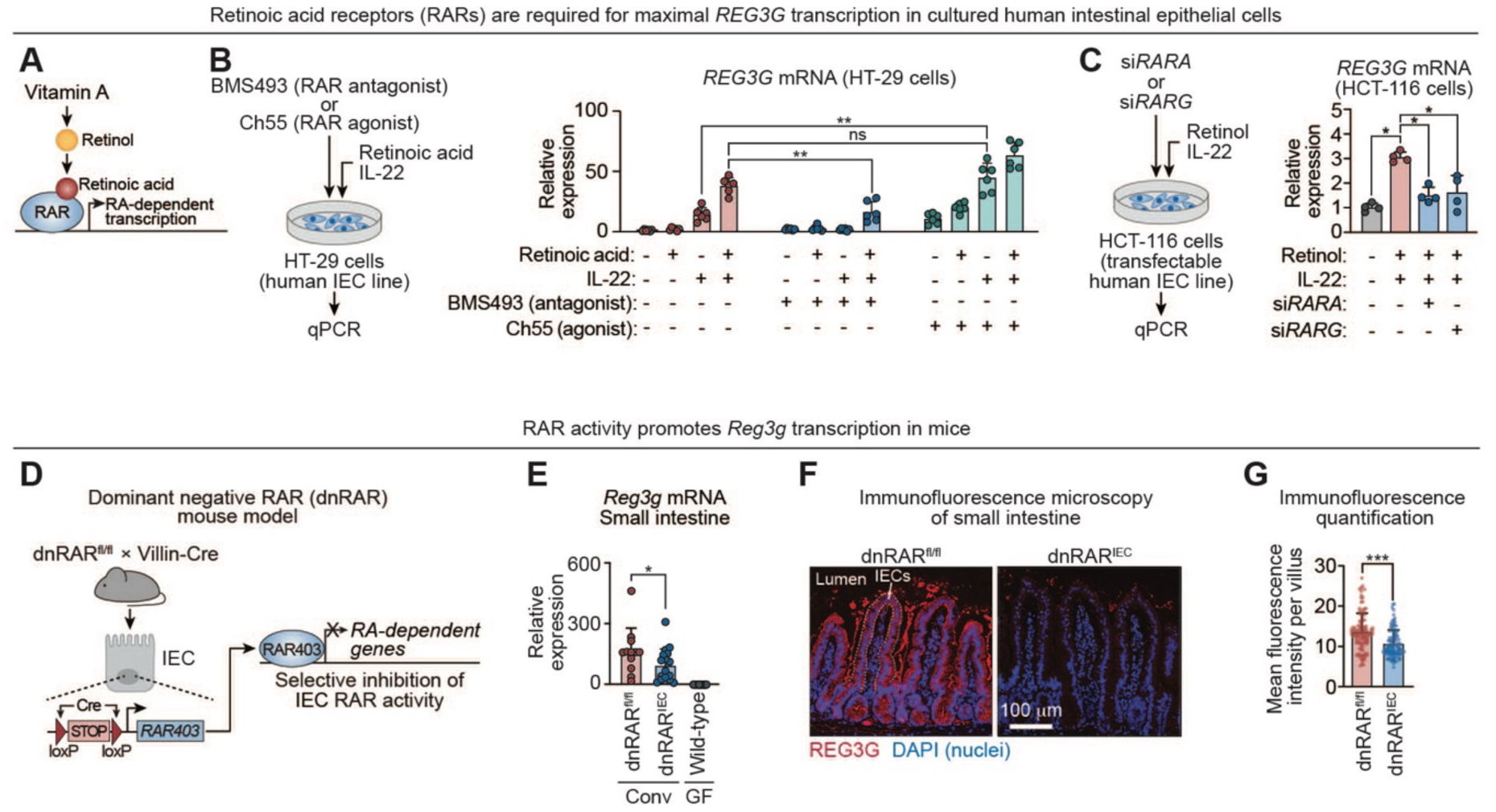
Retinoic acid receptors drive transcription of mouse *Reg3g* and human *REG3G* genes. **(A)** Vitamin A-derived retinol is metabolized to RA, which activates retinoic acid receptors (RARs). RARs bind target gene promoters to drive RA-dependent transcription. **(B)** HT-29 cells (human intestinal epithelial cells) were treated with the RAR antagonist BMS493 or the RAR agonist Ch55 and simultaneously stimulated overnight with RA and IL-22. *REG3G* transcripts were quantified by qPCR. Each data point represents an independent experimental replicate (n=6 per group). **(C)** HCT-116, a transfection-competent human intestinal epithelial cell line expressing *RARA* and *RARG* (Fig. S3B), was treated for 24 hours with an siRNA targeting either gene and then stimulated overnight with retinol and IL-22. *REG3G* transcripts were quantified by qPCR. Each data point represents an independent experimental replicate (n=4 per group). **(D)** IEC-specific disruption of RAR signaling using a dominant-negative RAR (dnRAR) knock-in allele. dnRAR mice harbor a *loxP*-flanked STOP cassette upstream of a dominant-negative RAR open reading frame. The dnRAR is derived from a mutant human RARα (RAR403) lacking the ligand-dependent transactivation domain and functions as a pan-RAR inhibitor. Crossing dnRAR mice with Villin-Cre transgenic mice excises the STOP cassette in IECs, resulting in IEC-selective expression of dnRAR and inhibition of RAR signaling. **(E)** qPCR analysis of *Reg3g* expression in small intestines of conventional dnRAR^fl/fl^ (n=12) and dnRAR^IEC^ (n=17) mice from five litters, and germ-free wild-type mice (n=21). **(F)** Immunofluorescence microscopy of REG3G in small intestines of dnRAR^fl/fl^ and dnRAR^IEC^ mice. Sections were stained for REG3G and counterstained with DAPI. Scale bar, 100 *μ*m. Images are representative of at least three fields per sample and two independent experiments (three littermates per group). **(G)** Mean fluorescence intensities of at least 150 villi from the images represented in (F) were quantified across at least two mice of each genotype. RAR, retinoic acid receptor; RA, retinoic acid; IEC, intestinal epithelial cell; REG3G, regenerating islet-derived protein 3γ; siRNA, small interfering RNA; dnRAR, dominant negative retinoic acid receptor; Conv, conventional; GF, germ-free. Means ± SEM are plotted; *p < 0.05; **p < 0.01; ***p<0.001; ns, not significant by Mann-Whitney test. See also Figure S3.

To complement these pharmacologic studies with a genetic approach, we performed siRNA-mediated knockdown of individual RARs. For these studies we used HCT-116 cells, a transfection-competent human intestinal epithelial cell line that, like HT-29 cells, upregulates *REG3G* transcription in response to IL-22 and retinol (Fig. S3A). HCT-116 cells express *RARA* and *RARG*, but not *RARB* (Fig. S3B). Knockdown of either *RARA* or *RARG* reduced *REG3G* transcript levels following stimulation with retinol and IL-22 (Fig. 3C; Fig. S3C). These findings further support a requirement for RARs in promoting *REG3G* transcription and suggest functional redundancy among RAR isoforms in human intestinal epithelial cells.

We next asked whether RAR activity similarly regulates mouse *Reg3g* expression in vivo. To address this, we used mice with intestinal epithelial cell–specific inhibition of RAR signaling. These mice harbor a knock-in allele containing *loxP*-flanked STOP sequences upstream of a dominant-negative human RARα (RAR403; “dnRAR”), which lacks the ligand-dependent transactivation domain and functions as a pan-RAR inhibitor (Fig. 3D).^34,35^ Crossing these mice with Villin-Cre transgenic mice results in epithelial cell-specific expression of dnRAR (dnRAR^IEC^) and suppression of RAR activity.^29^ Compared with dnRAR^fl/fl^ littermate controls, dnRAR^IEC^ mice showed reduced *Reg3g* transcription in the small intestine and decreased REG3G protein abundance by immunofluorescence microscopy (Fig. 3E–G; Fig. S3D). This reduction was not attributable to altered IL-22 availability, as *Il22* transcript levels were comparable between dnRAR^IEC^ mice and controls (Fig. S3E). Thus, epithelial RAR signaling is required to drive *Reg3g* expression in vivo.

### Retinoic acid receptors bind promoter RAREs to activate mouse *Reg3g* and human *REG3G* transcription

RARs canonically regulate gene expression by binding RAREs within target gene promoters (Fig. 4A).^32^ Because RA induces *Reg3g* expression in intestinal epithelial cells in an RAR-dependent manner, we asked whether

**Figure 4.**
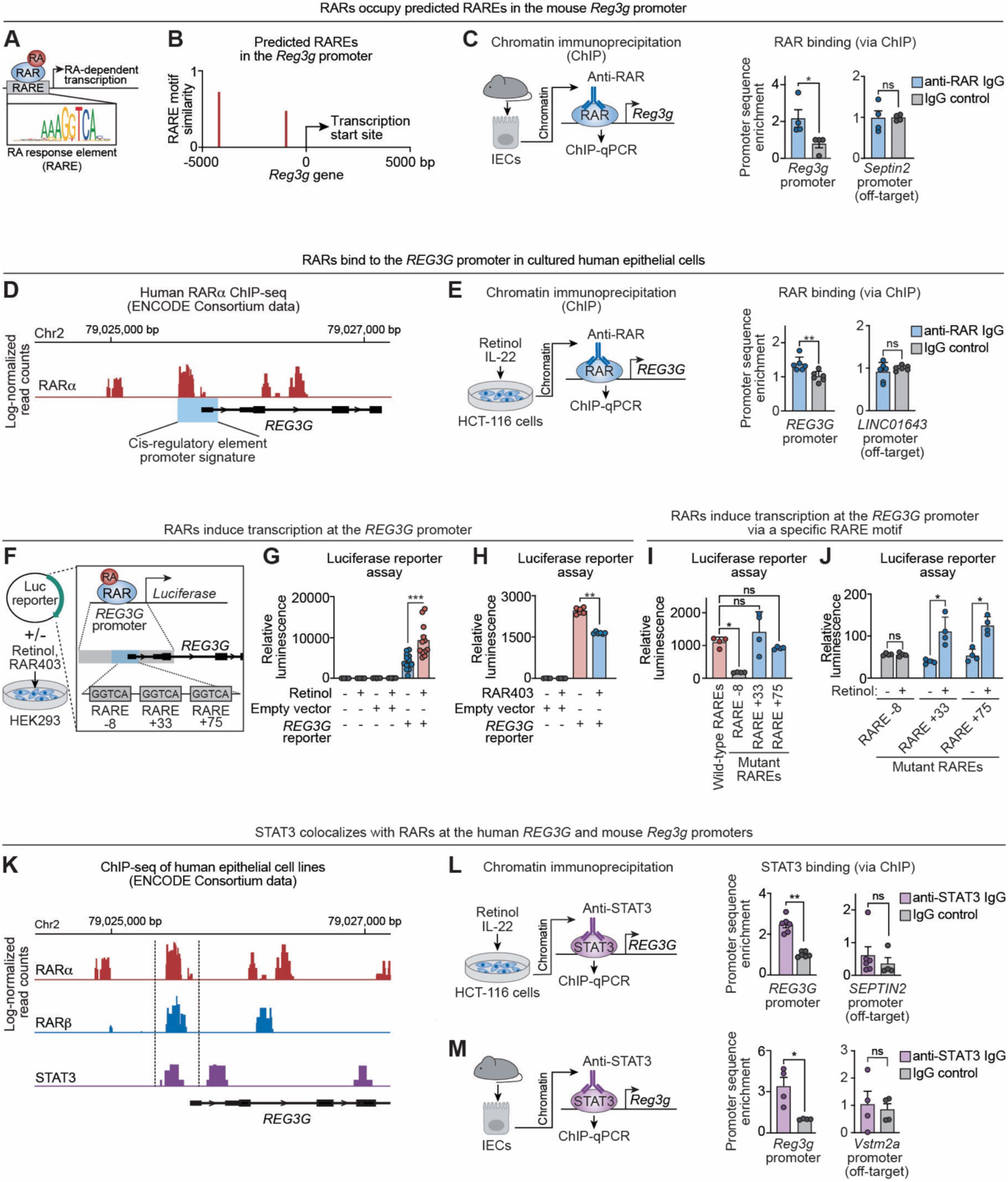
Retinoic acid receptors bind promoter RAREs to activate mouse *Reg3g* and human *REG3G* transcription. **(A)** RARs bind to retinoic acid response elements (RAREs) in target gene promoters to induce RA-dependent transcription. **(B)** The mouse *Reg3g* promoter was scanned for putative RAREs using the JASPAR motif database and quantified with the Transcription Factor Binding Site R Package.^56^ **(C)** IECs were isolated from wild-type mice, and RAR-bound chromatin was immunoprecipitated. Enrichment of *Reg3g* promoter regions and an off-target control promoter (*Vstm2a*) was quantified by qPCR using primers centered on the RARE at -4195 bp upstream of the transcription start site. Control ChIP used a non-specific IgG isotype control antibody. The data were from two mice, with two to three technical replicates per mouse. **(D)** ChIP-sequencing read tracks from HepG2 cells (human hepatocarcinoma-derived cells) expressing FLAG-tagged RARα, obtained from a publicly available ENCODE Consortium dataset^36^. Base pair positions are shown relative to the *REG3G* transcription start site. ENCODE candidate cis-regulatory element promoter signature is denoted by a blue box. **(E)** ChIP was performed on HCT-116 cells treated overnight with retinol and IL-22. RARs were immunoprecipitated using a pan-RAR antibody, and associated DNA was quantified by qPCR using primers centered on the RARα binding peak encompassing -212 bp to 8 bp relative to the *REG3G* start site. Binding to an off-target promoter (*LINC01643*) was also assessed. Control ChIP was done using a non-specific IgG isotype control antibody. Each data point represents an independent experimental replicate (n=6 per group). **(F)** Transcription reporter assays. HEK293 cells were transfected with a plasmid containing the native *REG3G* promoter region (2 kb) cloned upstream of a firefly luciferase reporter. The human *REG3G* promoter contains three predicted RAREs (RARE -8, +33, and +75). The cells were treated with 1 *μ*M retinol and/or co-transfected with a plasmid that expresses RAR403, which suppresses RAR activity (see Fig. 3D). **(G)** HEK293 cells were transfected with the *REG3G* promoter–luciferase reporter plasmid to establish that luminescence is dependent on the presence of the plasmid and retinol. Luciferase luminescence was measured in duplicate and each data point represents the average of two readings from one tissue culture well. (n=8 experimental replicates per group). **(H)** HEK293 cells were transfected with the *REG3G* promoter–luciferase reporter plasmid in the presence of retinol and the presence or absence of RAR403. Luminescence was measured in duplicate and each data point represents average readings from one tissue culture well. (n=6 experimental replicates per group). **(I)** Luciferase reporter assay comparing transcriptional activity of the native human promoter and promoter variants with individual disruptions of the RAREs shown in (F). Assays were performed in the presence of retinol. Each data point represents an independent experimental replicate (n=4 per group). **(J)** Luciferase reporter assay comparing transcriptional activity of the native human promoter and promoter variants with individual RARE disruptions, assessed in the presence or absence of retinol. Each data point represents an independent experimental replicate (n=4 per group). **(K)** ChIP-sequencing read tracks from ENCODE datasets showing binding of tagged RARα, RARβ, and STAT3 upstream of and within the *REG3G* locus. Data were generated in human epithelial cells lines engineered to express FLAG-tagged RARα (HepG2), RARβ (A549), or endogenous STAT3 (MCF-10A-derived). **(L)** Chromatin was isolated from cultured HCT-116 cells and STAT3-bound chromatin was immunoprecipitated. Enrichment of *REG3G* promoter sequences and an off-target control promoter (*SEPTIN2*) were quantified by qPCR. Control ChIP used a non-specific IgG isotype control antibody. **(M)** Chromatin was isolated from IECs recovered from wild-type mice and STAT3-bound chromatin was immunoprecipitated. Enrichment of *Reg3g* promoter sequences (centered on RARE -4195) and an off-target control promoter (*Vstm2a*) were quantified by qPCR. Control ChIP used a non-specific IgG isotype control antibody. The data were generated from two mice, with two to three technical replicates per mouse. RA, retinoic acid; RAR, retinoic acid receptor; RARE, retinoic acid response element; REG3G, regenerating islet-derived protein 3γ; IEC, intestinal epithelial cell; ChIP, chromatin immunoprecipitation; qPCR, quantitative real-time PCR; Chr2, chromosome 2; Luc, luciferase; STAT3, Signal Transducer and Activator of Transcription 3. Means ± SEM are plotted; *p < 0.05; **p < 0.01; ***p <0.001; ns, not significant by Mann-Whitney test. See also Figure S4.

RARs bind at the *Reg3g* promoter in vivo. Motif scanning of the mouse *Reg3g* promoter identified two candidate RAREs, located 4195 bp and 970 bp upstream of the transcription start site (Fig. 4B). Chromatin immunoprecipitation (ChIP) of isolated mouse intestinal epithelial cells revealed enrichment of promoter sequences encompassing the -4195 bp site in anti-RAR immunoprecipitates relative to IgG controls, whereas no enrichment was detected at an off-target promoter (Fig. 4C). These data indicate that RARs bind at the mouse *Reg3g* promoter in vivo.

We next examined RAR binding at the human REG3G promoter. Analysis of publicly available ENCODE ChIP-sequencing datasets^36^ and ENCODE promoter annotations^37^ revealed RARα occupancy upstream of the *REG3G* transcription start site and within its predicted promoter region (Fig. 4D; Fig. S4A). We validated this experimentally by performing ChIP in HCT-116 cells treated with retinol and IL-22. Anti-RAR immunoprecipitates were enriched for *REG3G* promoter sequences relative to IgG controls, whereas no enrichment was observed at an off-target promoter (Fig. 4E). Thus, RARs bind at both mouse *Reg3g* and human *REG3G* promoters.

To determine whether RAR binding drives transcription of *REG3G*, we performed luciferase reporter assays in HEK293 cells. Cells were transfected with reporter constructs carrying either a synthetic RARE–driven promoter or a 2-kb fragment of the native human *REG3G* promoter encompassing the RAR binding region identified by ENCODE analysis (Fig. 4D, F; Fig. S4B). Retinol increased luciferase activity from both reporters, whereas co-expression of the dominant-negative RAR (RAR403) impeded this induction (Fig. 4G, H; Fig. S4C, D). These results indicate that retinoid-driven activation of the *REG3G* promoter requires functional RAR signaling.

We next defined the specific RAREs required for promoter activation. Fine mapping identified three candidate RARE motifs at -8, +33, and +75 bp relative to the *REG3G* transcription start site (Fig. 4F). Mutation of RARE -8 reduced retinol-dependent luciferase activity, whereas mutation of RARE +33 or RARE +75 had no statistically significant effect (Fig. 4I, J). Thus, RARE -8 is the primary functional element mediating RAR-dependent activation of the *REG3G* promoter.

Microbial signals promote *Reg3g* expression through IL-22–dependent activation of STAT3.^7–9^ Reanalysis of ENCODE datasets revealed co-occupancy of STAT3 with RARα and RARβ at the human *REG3G* promoter (Fig. 4K). Consistent with this, ChIP analysis of HCT116 cells and mouse intestinal epithelial cells revealed STAT3 binding adjacent to RAR-occupied regions of the *Reg3g/REG3G* promoters (Fig. 4L, M). Together, these findings indicate that nutritional and microbial cues converge at the *Reg3g*/*REG3G* promoters through adjacent binding of RARs and STAT3, thereby integrating RA and IL-22 signaling to drive antimicrobial gene expression.

### Vitamin A enhances expression of α-defensin antimicrobial proteins in the small intestine

Antimicrobial defense in the small intestine is mediated by multiple epithelial protein families with distinct structures and mechanisms of action. In addition to REG3 lectins, Paneth cells produce α-defensins, a family of cationic antimicrobial peptides that, like REG3 lectins, kill bacteria by disrupting their membranes (Fig. 5A).^38^ Unlike *Reg3* genes, α-defensin gene expression in mice is largely independent of microbial colonization, making this family an ideal system to test whether vitamin A broadly regulates epithelial antimicrobial programs rather than selectively amplifying microbiota-responsive genes.

**Figure 5.**
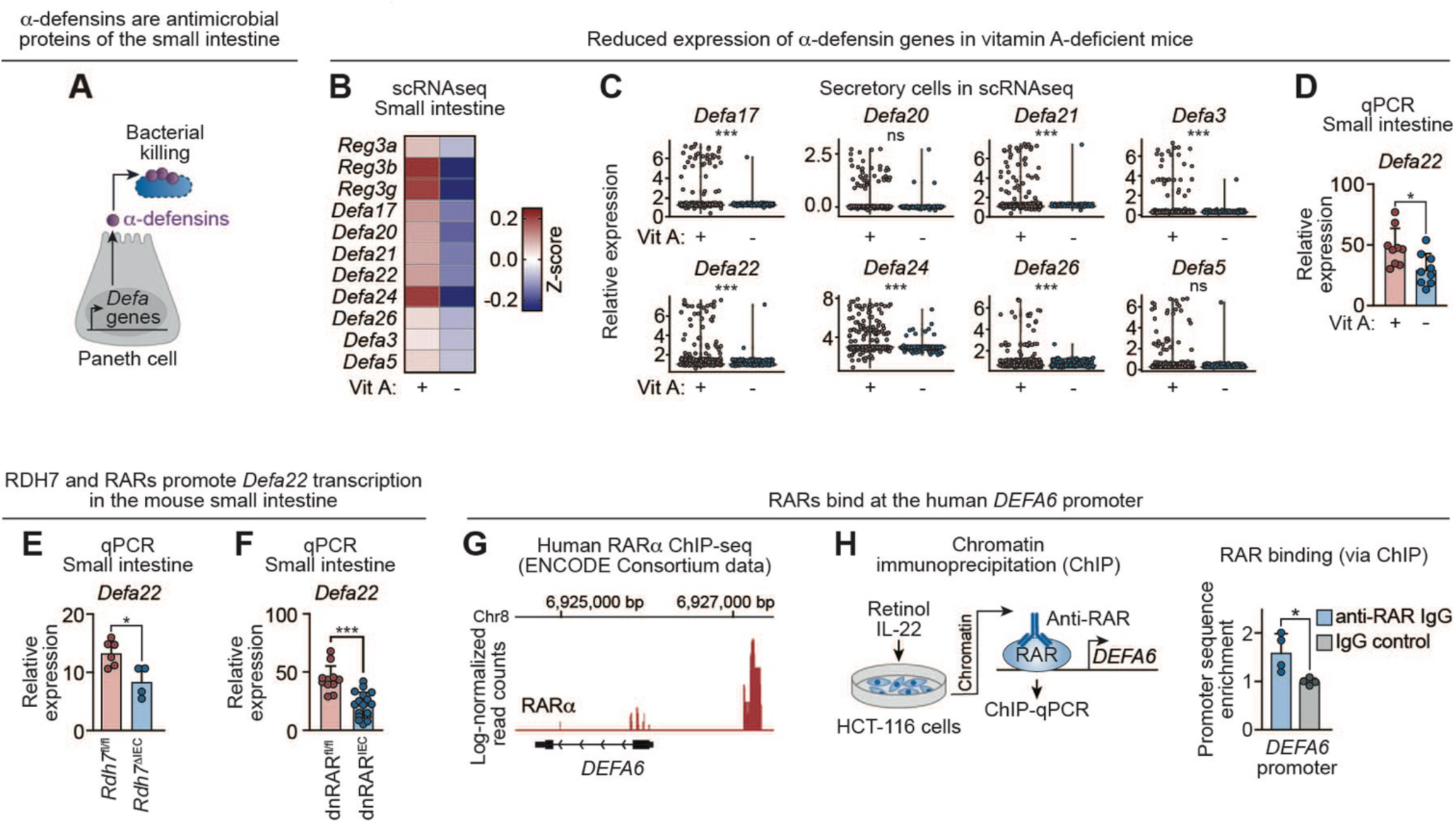
Vitamin A enhances expression of α-defensin antimicrobial proteins in the small intestine. **(A)** α-defensins are a family of antibacterial proteins produced by Paneth cells. In contrast to REG proteins, α-defensin gene expression in mice is not induced by the microbiota. **(B)** Heatmap of averaged Z-scores of antimicrobial genes from scRNA-seq data of epithelial cells isolated from Vit A+ and Vit A– mice as described in Fig. 1A. **(C)** Violin plots of α-defensin gene expression in secretory cells from the scRNA-seq dataset shown in Fig. 1. Each data point represents an individual cell. P values from DESeq2 comparative analysis are shown above each plot. **(D)** qPCR analysis of *Defa22* transcripts in the small intestine of Vit A+ (n=9) and Vit A– (n=9) mice. Each data point represents one mouse. **(E)** qPCR analysis of *Defa22* transcripts in the small intestines of conventional *Rdh7*^fl/fl^ (n=6) and *Rdh7*^μIEC^ (n=4) littermates (from two litters). Each data point represents one mouse. **(F)** qPCR analysis of *Defa22* transcripts in the small intestines of conventional dnRAR^fl/fl^ (n=12) and dnRAR^IEC^ (n=17) littermates from five litters. Each data point represents one mouse. **(G)** Chromatin immunoprecipitation (ChIP)-sequencing read tracks from HepG2 cells (human hepatocarcinoma-derived cells) expressing FLAG-tagged RARα, obtained from a publicly available ENCODE Consortium dataset^36^. Base pair positions are shown relative to the *DEFA6* transcription start site. **(H)** IECs were isolated from wild-type mice and RAR-bound chromatin was immunoprecipitated. Enrichment of *DEFA6* promoter sequences (encompassing -1273 to -1070 bp upstream of the *DEFA6* transcription start site) was quantified by qPCR. Control ChIP was performed using a non-specific IgG isotype control antibody. Data were generated from two mice, with two technical replicates per mouse. DEFA, α-defensin; scRNA-seq, single cell RNA sequencing; qPCR, quantitative real-time PCR; Vit A, vitamin A; RAR, retinoic acid receptor; dnRAR, dominant negative retinoic acid receptor; ChIP, chromatin immunoprecipitation. Means ± SEM are shown; *p < 0.05; ***p<0.001; ns, not significant by Mann-Whitney test unless otherwise specified.

To address this question, we revisited our single-cell RNA-sequencing dataset from intestinal epithelial cells isolated from vitamin A–replete and vitamin A–deficient mice. This analysis revealed a coordinated reduction in transcripts encoding multiple α-defensin genes (*Defa* family) in vitamin A–deficient animals (Fig. 5B). Within the Paneth cell compartment, this reduction reflected both decreased transcript abundance and a lower fraction of cells expressing individual *Defa* paralogs, including *Defa17*, *Defa21*, *Defa22*, *Defa24*, and *Defa26* (Fig. 5C).

We focused subsequent validation experiments on *Defa22* as a representative α-defensin gene. Quantitative PCR confirmed reduced *Defa22* transcript levels in the small intestines of vitamin A–deficient mice compared with vitamin A–replete controls (Fig. 5D). *Defa22* expression was similarly diminished in *Rdh7*^μIEC^ mice relative to *Rdh7*^fl/fl^ littermates, indicating a requirement for epithelial conversion of retinol to RA (Fig. 5E). Likewise, *Defa22* expression was reduced in dnRAR^IEC^ mice compared with dnRAR^fl/fl^ controls, demonstrating dependence on RAR signaling (Fig. 5F). These findings establish that α-defensin genes, like *Reg3* genes, are regulated by vitamin A through RA–dependent activation of RARs.

To determine whether this regulatory mechanism is conserved in humans, we analyzed publicly available ENCODE ChIP -sequencing data^36^ and identified a prominent RARα binding site approximately 1 kb upstream of the human *DEFA6* transcription start site (Fig. 5G). ChIP–qPCR in HCT-116 cells confirmed RAR binding at this promoter region (Fig. 5H). Together, these data identify a shared vitamin A–RAR regulatory axis that promotes epithelial antimicrobial programs across multiple gene families, including REG3 lectins and α-defensins.

## Discussion

The intestinal epithelial surface is in direct contact with dense microbial communities and must rapidly deploy innate defense mechanisms to preserve barrier integrity. Although vitamin A is well-established as a regulator of intestinal adaptive immunity, its role in epithelial innate immunity has been less clear. Here, we identify vitamin A as a direct regulator of epithelial antimicrobial programs in the gut. We show that epithelial conversion of retinol to RA activates RARs that drive transcription of genes encoding key antimicrobial proteins. This epithelial cell–intrinsic pathway provides a mechanism by which nutritional status directly shapes first line mucosal defense. These findings expand the immunological functions of vitamin A beyond adaptive immunity and identify it as a modulator of rapid innate responses at the intestinal barrier.

Microbial induction of intestinal *Reg3g* expression occurs through a well-defined multicellular signaling cascade. In this pathway, microbial ligands stimulate myeloid cells to produce IL-23, leading to IL-22 secretion by group 3 innate lymphoid cells and subsequent STAT3 activation in epithelial cells.^5,6,8,9^ Our results define a parallel, epithelial-intrinsic pathway in which vitamin A metabolism amplifies antimicrobial gene expression through RAR signaling. Whereas microbial cues establish baseline *REG3G* expression, vitamin A functions as a second signal that enhances transcriptional output. At the molecular level, RARs and STAT3 bind adjacent regions of the *Reg3g*/*REG3G* promoters in mice and humans, identifying a genomic locus where dietary and microbial signals converge. This promoter-level integration provides a mechanistic framework for how epithelial cells jointly interpret nutritional and microbial inputs to regulate antimicrobial defense.

Our findings further identify epithelial RARs as coordinators of both innate and adaptive immunity. We previously showed that epithelial RAR signaling drives expression of serum amyloid A (SAA) proteins—retinol carriers induced by the microbiota through STAT3 and likewise under RAR control.^13,29^ Thus, SAAs and antimicrobial proteins share a regulatory architecture in which microbial and vitamin A–dependent signals converge at the level of transcription. Functionally, however, these two outputs are distinct. Antimicrobial proteins act locally and rapidly at the epithelial surface to limit microbial encroachment.^25,26^ In contrast, SAAs export retinol from epithelial cells to lamina propria myeloid cells, enabling RA production that supports T- and B-cell development.^13^ Together, these findings suggest that a common STAT3–RAR module coordinates two temporally and spatially distinct programs: an immediate epithelial antimicrobial response and a downstream adaptive immune response. Through this shared transcriptional circuitry, epithelial cells couple barrier defense to the programming of longer-term immunity.

The requirement for vitamin A in regulating intestinal antimicrobial programs parallels findings in the skin. In the mouse epidermis, expression of the antimicrobial protein RELMα — the only previously described vitamin A–regulated antimicrobial protein — is similarly enhanced by vitamin A signaling.^39^ More broadly, these observations support a recurring role for lipid-soluble vitamins as transcriptional regulators of epithelial antimicrobial programs. Vitamin D provides a striking example: it promotes transcription of the antimicrobial peptide cathelicidin in keratinocytes through activation of the vitamin D receptor, which binds regulatory elements in the cathelicidin gene promoter.^40,41^ Together, these studies suggest a general principle in which lipophilic vitamins link micronutrient availability to barrier immunity across epithelial tissues.

Why might vitamin A control antimicrobial protein expression? One possibility is that vitamin A serves as a signal of nutritional sufficiency. Because its absorption depends on adequate dietary fat,^42^ vitamin A availability could reflect sufficient metabolic resources to support the energetically demanding synthesis and secretion of antimicrobial proteins. In this view, vitamin A functions as a contextual cue aligning antimicrobial output with nutritional state.

A second possibility is that vitamin A helps to protect host tissues from the potential toxicity of antimicrobial proteins. REG3 lectins and α-defensins disrupt bacterial membranes but can also damage host membranes at high local concentrations.^43,44^ Vitamin A–derived metabolites may alter epithelial membrane properties in ways that confer resistance to collateral injury during antimicrobial deployment. An analogous strategy is used by *Staphylococcus aureus*, which incorporates carotenoid pigments—structurally related to vitamin A—into its membranes to increase resistance to membrane-disrupting peptides.^45^ Although speculative, this model raises the possibility that vitamin A coordinates antimicrobial deployment with epithelial self-protection.

An apparent paradox arising from our findings is that expression of RDH7, the epithelial enzyme required for conversion of retinol to RA, is suppressed by the gut microbiota.^30^ At first glance, this seems counterintuitive, as microbial exposure might be expected to enhance RDH7 expression to promote antimicrobial gene induction. One explanation is that activation of antimicrobial programs requires only limited or transient retinol metabolism. Following this initial phase, retinol may be redirected away from epithelial retinoic acid production and toward other immunological functions. In light of our prior work showing that epithelial cells export retinol via SAA to support adaptive immune programming, microbiota-dependent downregulation of RDH7 may reflect a coordinated shift in retinoid allocation. In this model, early epithelial RA production promotes rapid barrier defense, whereas subsequent redistribution of retinol to immune cells supports longer-term adaptive immunity. Thus, retinoid metabolism may be dynamically partitioned across immune compartments over time, aligning micronutrient utilization with the evolving phases of host defense.

Several limitations of this study warrant consideration. First, although our dietary and culture-based depletion strategies demonstrate that vitamin A regulates *Reg3g/REG3G* expression, these approaches do not reduce expression to germ-free levels. However, complete vitamin A deprivation in vivo is precluded due to maternal hepatic stores and transfer through breastmilk^46,47^, resulting in residual vitamin A availability in offspring.^48,49^ Similar constraints apply to epithelial cell culture systems, where prolonged vitamin A deprivation compromises viability.^50,51^ Thus, whether vitamin A serves as an absolute requirement or instead tunes the magnitude of antimicrobial gene induction remains unresolved.

Second, while we establish a requirement for RAR signaling, the contributions of individual RAR isoforms remain unresolved. Our dnRAR model establishes that RAR-dependent transcription is essential but does not distinguish among RAR paralogs. In human intestinal epithelial cells, knockdown of either *RARA* or *RARG* reduced *REG3G* expression, consistent with functional redundancy. A similar redundancy may operate in the in vivo intestinal epithelium, where multiple epithelial lineages contribute to antimicrobial defense. Isoform-specific effects are also likely to depend on cellular context. For example, epithelial-specific deletion of *Rara* produces broad alterations in epithelial gene expression and homeostasis, highlighting the context-dependent nature of RAR function.^52^ In addition, interactions with corepressors and coactivators, including NCOR2/SMRT, may further shape transcriptional output.^53^ Thus, while our data define an essential role for RAR signaling in regulating epithelial antimicrobial programs, delineating isoform-specific contributions will require future studies in defined epithelial contexts.

Finally, although we show adjacent binding of RARs and STAT3 at the *Reg3g*/*REG3G* promoters, we have not determined whether these factors physically interact or cooperatively remodel chromatin at these loci. Defining the molecular architecture of this transcriptional integration will further clarify how dietary and microbial signals are mechanistically coupled.

In summary, this study identifies a previously unrecognized role for vitamin A in regulating intestinal innate immunity through epithelial cell-intrinsic control of antimicrobial protein expression. Coordinated binding of RARs and STAT3 at the *Reg3g/REG3G* promoters provides a mechanistic framework for integration of dietary and microbial signals. By extending this RAR-dependent regulatory axis to multiple antimicrobial protein families, including α-defensins, our findings establish vitamin A as a key nutritional regulator of epithelial antimicrobial programs. These results help explain the long-recognized association between vitamin A status and susceptibility to intestinal infection^54,55^ and highlight nutritional regulation of epithelial immunity as a central determinant of host–microbe interactions at barrier surfaces.

## Resource Availability

Raw sequencing files are available at PRJNA1416395. Processed single cell RNAseq data is available at GSE318054 and at the Single Cell Portal SCP3497.

## Contact for Reagent and Resource Sharing

Further information and requests for resources and reagents should be directed to and will be fulfilled by the Lead Contact, Lora V. Hooper.

## Acknowledgments

We thank J. Shelton at the UT Southwestern Histology Core for assistance with preparing tissue slides, C. Arana and B. Zhang at the UT Southwestern Microbiome Research Laboratory for assistance with single cell RNA library preparation and sequencing, Dr. Shanthini Sockanathan (Johns Hopkins University) for providing the dnRAR^fl/fl^ mice, and the BioHPC at UT Southwestern for their support of GPU systems and coding guidance. This work was supported by NIH grants R01 DK070855 (L.V.H.) and R01 HL151650 (N.V.M.), Welch Foundation Grant I-1874 (L.V.H.), the Walter M. and Helen D. Bader Center for Research on Arthritis and Autoimmune Diseases (L.V.H.), and the Howard Hughes Medical Institute (L.V.H.). G.Q. was supported by NIH T32 AI005284. Components of some figures were generated using BioRender.

## Author Contributions

G.Q., J.G.M., N.V.M. and L.V.H designed the research. G.Q., D.C.P., K.K., M.J., G.V., A.J., B.H., and C.L.B. performed the research. G.Q., D.C.P., G.V., A.J., analyzed data. G.Q. and L.V.H. wrote the paper.

## Declaration of generative AI and AI-assisted technologies in the manuscript preparation process

We used OpenAI’s ChatGPT (version 4.0, 5.0, and 5.2) solely for language editing (e.g., grammar, clarity, and brevity) and for suggesting alternative phrasings during manuscript preparation. The tool did not contribute to study design, data generation, and analysis. It was not used to generate references. All suggestions from the ChatGPT were reviewed, revised, and approved by the authors. No text was inserted verbatim without subsequent human editing. The authors take full responsibility for the content of the manuscript.

---

**Figure S1.**
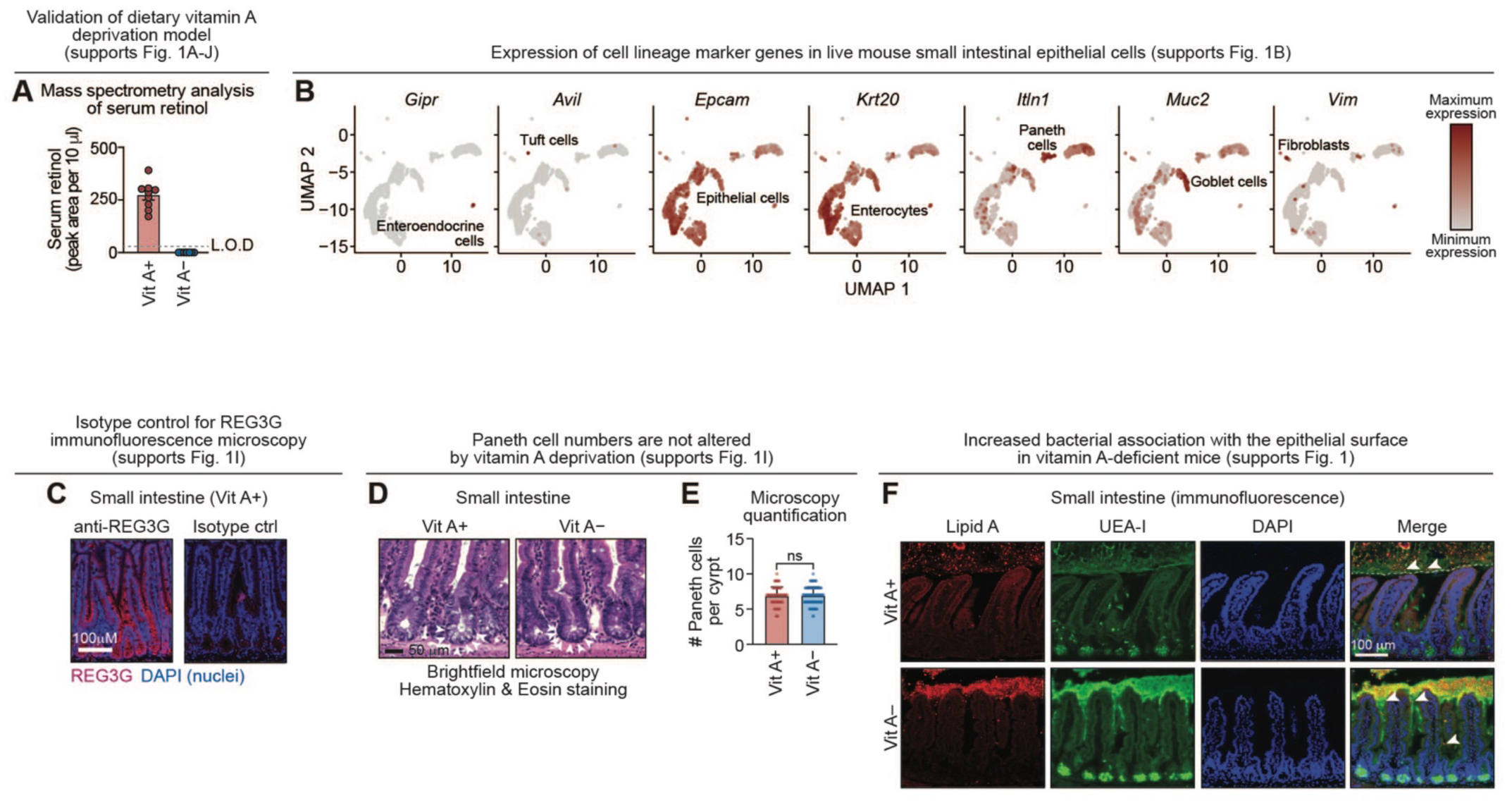
Characterization of the vitamin A deprivation model and expression of epithelial cell marker genes as determined by scRNAseq (supports Figure 1). **(A)** Mass spectrometry measurement of retinol from serum of mice fed provided a Vit A+ (n=9) or Vit A– (n=14) diet as described in Fig. 1A. Each dot represents one mouse, and the limit of detection (L.O.D.) is indicated. **(B)** Expression of marker genes among small intestinal cell populations as determined by scRNA-seq. **(C)** Immunofluorescence microscopy of REG3G in small intestine sections from Vit A+ mice, with comparison to an isotype control antibody. Sections were stained for REG3G (red) and counterstained with DAPI (blue) to detect nuclei. Scale bar, 100 *μ*m. **(D)** Representative images of small intestinal crypts of mice fed a Vit A+ or Vit A– diet. Paneth cells were identified by their distinctive morphology and the presence of dense secretory granules (white arrowheads). Scale bar, 50 *μ*m. **(E)** Enumeration of Paneth cells identified in crypts from Vit A+ (60 crypts counted across 5 mice) and Vit A–mice (94 crypts counted across four mice). **(F)** Immunofluorescence microscopy of lipid A (bacteria) and UEA-I (mucus) in small intestine sections from Vit A+ mice and Vit A- mice. The tissue was counterstained with DAPI to detect nuclei. The mucus barrier is outlined with a white dotted line and examples of positive lipid A staining are indicated by white arrow heads. Vit A, vitamin A; L.O.D., limit of detection; REG3G, regenerating islet-derived protein 3γ; Ctrl, control; UEA-I, Ulex Europaeus Agglutinin I. Means ± SEM are plotted; ns, not significant by Mann-Whitney test.

**Figure S2.**
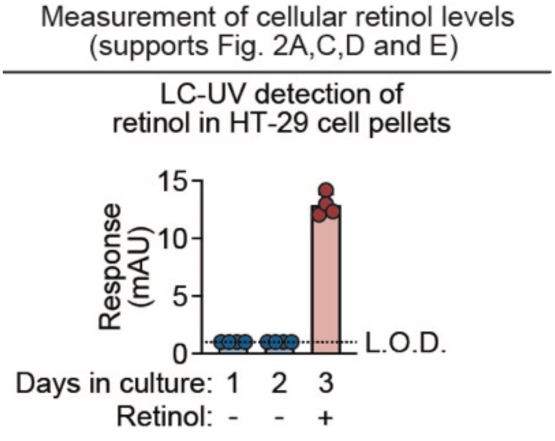
Characterization of retinol levels in HT-29 cells (supports Figure 2). Liquid chromatography–UV spectroscopy analysis of cell pellets from HT-29 cells grown in DMEM with 10% charcoal stripped fetal bovine serum for three consecutive days. On day 3, 1 *μ*M retinol was added to the culture medium and the cells were incubated for an additional 12 hours before analysis. Each data point represents one experimental replicate (n=4 per group). LC–UV, liquid chromatography ultraviolet spectroscopy; L.O.D., limit of detection; mAU (milli absorbance units). Means are plotted.

**Figure S3.**
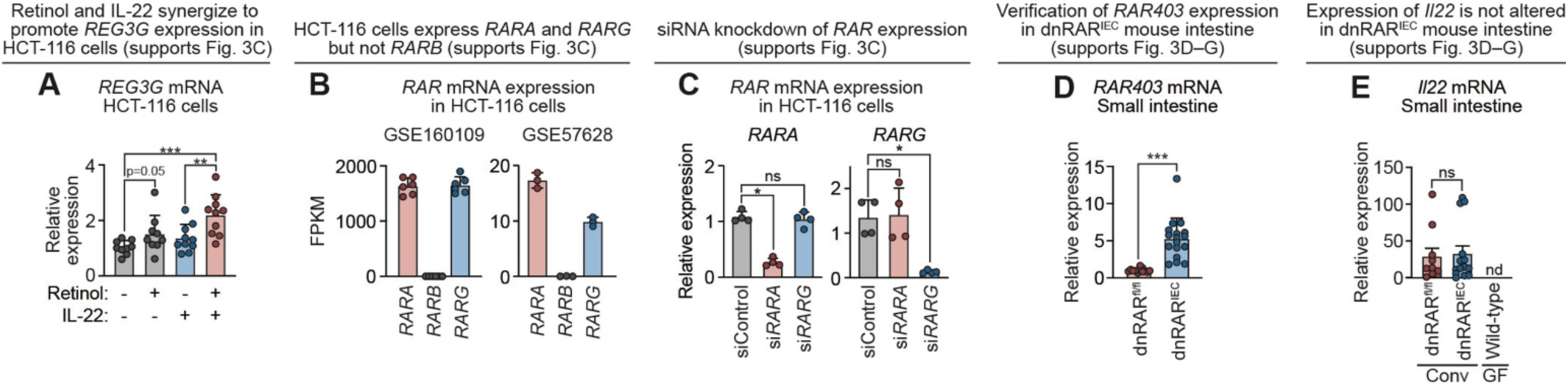
Characterization of *REG3G* and *RAR* expression in HCT-116 cells and validation of a genetic mouse model of RAR deficiency (supports Figure 3). **(A)** qPCR analysis of *REG3G* transcripts in HCT-116 cells treated with or without retinol and/or IL-22. Each data point represents one experimental replicate (n=9 or 10 replicates per group). **(B)** FPKM values of all three RAR gene paralogs (*RARA*, *RARB*, *RARG*) in HCT-116 cells. Data are from two independent and previously published RNA-sequencing studies (GSE160109 and GSE57628).^57^ Each data point represents one experimental replicate (n=6 for GSE160109 and n=3 for GSE57628). **(C)** qPCR of *RARA* and *RARG* expression 48 hours after siRNA treatment. Each data point represents one experimental replicate (n=4 per group). **(D)** qPCR of *RAR403* transcripts in small intestines of conventional dnRAR^fl/fl^ (n=12) and dnRAR^IEC^ (n=17) littermates (from five litters). Each data point represents one mouse. **(E)** qPCR of Il22 transcripts in small intestines of conventional dnRAR^fl/fl^ (n=10) and dnRAR^IEC^ (n=14) littermates (from five litters) and germ-free wild-type mice (n=8). Each data point represents one mouse. REG3G, regenerating islet-derived protein 3γ; RAR, retinoic acid receptor; siRNA, small interfering RNA; dnRAR, dominant negative retinoic acid receptor; Conv, conventional; GF, germ-free; nd, not detectable. All qPCR measurements were performed in at least duplicate. Each data point represents one tissue culture well or one mouse where applicable. Means ± SEM are plotted; *p < 0.05; ***p<0.001; ns, not significant by Mann-Whitney test.

**Figure S4.**
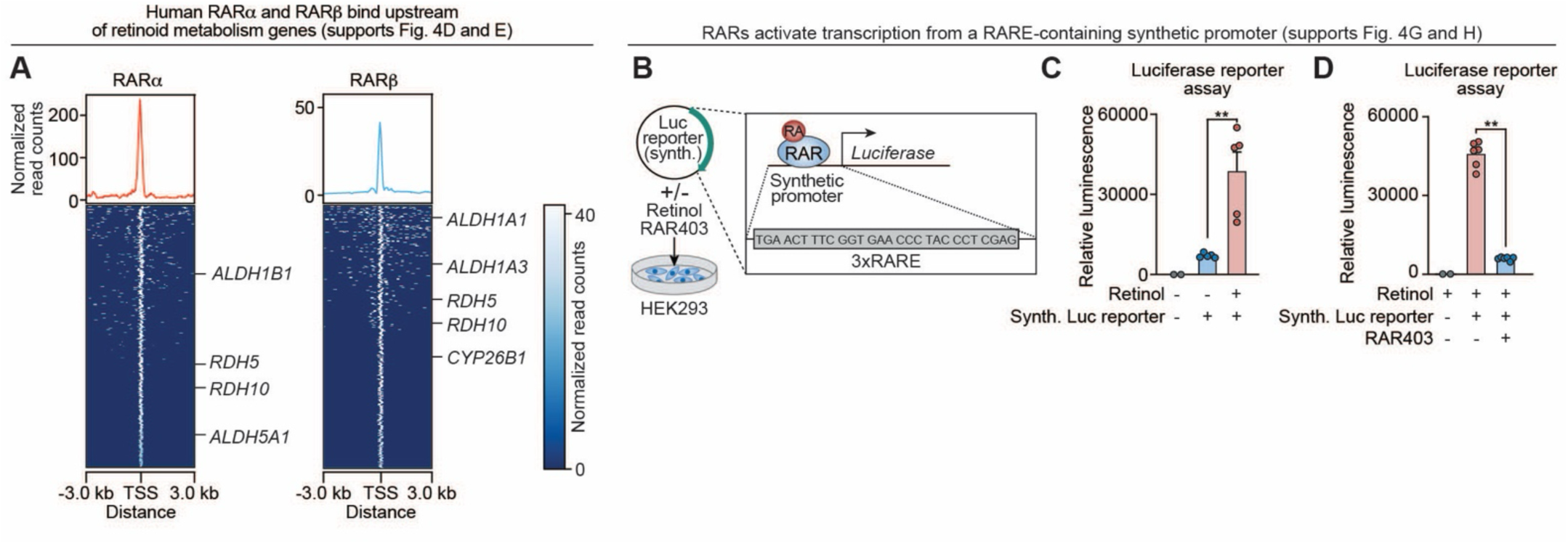
Controls for binding of RARα and RARβ in the human genome and characterization of transcription reporter assays (supports Figure 4). **(A)** ChIP sequencing heatmap peaks centered around transcription start sites (TSS) generated via Deeptools^58^. Rows corresponding to retinoid metabolism genes are labeled. **(B)** Transcription reporter assay with a synthetic RARE-containing promoter. HEK293 cells were transfected with a pGL2 plasmid containing a synthetic promoter sequence containing eight canonical RAREs positioned directly upstream of the firefly luciferase gene (synth. Luc reporter). The cells were treated with 1 *μ*M retinol and/or co-transfected with a plasmid that expresses RAR403, which impairs RAR activity (see Fig. 3D). **(C)** Luciferase reporter assay in the presence or absence of 1 *μ*M retinol. HEK293 cells were transfected with the synthetic promoter–Luciferase reporter plasmid to establish that luminescence is dependent on the presence of the plasmid. Luciferase luminescence was measured in duplicate and each data point represents the average of two readings from one tissue culture well. (n=5 experimental replicates per group). **(D)** HEK293 cells were transfected with the synthetic promoter–Luciferase reporter plasmid in the presence of 1 *μ*M retinol and with and without RAR403. Luminescence was measured in duplicate and each data point represents the average readings from one tissue culture well. (n=6 experimental replicates per group). RAR, retinoic acid receptor; TSS, transcription start site; Luc, luciferase; RA, retinoic acid; RARE, retinoic acid response element; Synth, synthetic. All luciferase measurements were performed in at least duplicate. Each bar graph data point represents one tissue culture well where applicable. Means ± SEM are plotted; **p < 0.01 by Mann-Whitney test.

## Methods

### Mice

Wild-type C57BL/6 mice were purchased from Jackson Laboratory. *Rdh7^-/-^* and *Rdh7*^fl/fl^ mice were generated in the UT Southwestern transgenic core and validated as previously described.^30^ Homozygous *Rdh7*^fl/fl^ mice were crossed with Villin-Cre transgenic mice (B6.Cg-Tg(Vil1-cre)997Gum/J; Jackson Laboratory)^59^ to generate intestinal epithelial cell-specific knockouts. Mice carrying a floxed dominant-negative allele of human *RARA* (dnRAR^fl/fl^)^35^ were provided by Dr. Shanthini Sockanathan (Johns Hopkins University) and crossed with Villin-Cre transgenic mice to generate dnRAR^IEC^ animals. All mouse strains were bred and maintained under specific pathogen-free conditions at the University of Texas Southwestern Medical Center.

Mice were fed ad libitum with standard chow (Lab Diet 6F5KAI), a vitamin A–deficient diet (Inotiv TD.09838; Envigo) or the matched vitamin A–replete diet (Inotiv TD.09830; Envigo). Germ-free C57BL/6 mice were bred and maintained in isolators at UT Southwestern. Both male and female mice were used and were analyzed between 7 and 20 weeks of age. To standardize microbial exposure prior to experimentation, all mice received either an oral gavage of segmentous filamentous bacteria (SFB)–positive fecal material (confirmed by PCR using eubacterial primers; Table S1) and/or a bedding transfer from C57BL/6 mice purchased from Taconic Farms. All animal experiments were approved by the UT Southwestern Institutional Animal Care and Use Committee.

### Dietary vitamin A deprivation of mice

Timed-pregnant C57BL/6 dams were maintained on either a vitamin A–replete diet (TD.09839; Envigo), containing 20,000 IU/kg retinyl acetate, or a nutritionally-matched vitamin A–deficient diet (TD.09838; Envigo) beginning at embryonic day 9. Pups were weaned at postnatal day 21 and maintained on the same dietary regimen until analysis.^12,23,60^ Animals were sacrificed between 8 and 12 weeks of age, and tissues were recovered for downstream analyses.

### Intraperitoneal injection of retinoic acid

Mice received two intraperitoneal injections of 100 *μ*L of freshly prepared all-trans retinoic acid (Sigma-Aldrich) dissolved in dimethyl sulfoxide (DMSO) at a concentration of 25 *μ*g/*μ*L. Injections were administered 24 hours and 8 hours prior to sacrifice. Control mice received two injections of vehicle alone (100 *μ*L DMSO) at the same time points.

### Cell culture

HT-29, HCT-116, and HEK293 cell lines (ATCC) were cultured in 1X DMEM supplemented with GlutaMAX (Gibco), penicillin-streptomycin, and 10% fetal bovine serum (FBS) at 37°C in a humidified incubator with 5% CO₂. Cells were passaged using 0.25% trypsin-EDTA and seeded at assay-appropriate densities. Prior to retinoid or cytokine treatment, cells were cultured for 24 hours in DMEM containing 10% charcoal-stripped FBS to deplete background retinoids.

### Retinoid, cytokine, and pharmacologic treatment of cultured cells

Retinol and all-trans retinoic acid (Sigma-Aldrich) were dissolved in DMSO under light-protected conditions and used at a final concentration of 1 µM. Vehicle controls received equivalent volumes of DMSO. Recombinant human IL-22 (Abcam) was added at a final concentration of 100 ng/mL and incubated overnight (16–18 hours), either alone or in combination with retinol or retinoic acid.

Where indicated, retinol metabolism was inhibited using disulfiram (Tocris), added at a final concentration of 100 µM concurrently with retinol and IL-22. For rescue experiments, retinoic acid was added simultaneously with disulfiram and other treatments.

Pharmacologic modulation of retinoic acid receptor (RAR) activity was performed using the pan-RAR antagonist BMS493 (Fisher Scientific) or the pan-RAR agonist Ch55 (Tocris). These compounds were added concurrently with retinoic acid and IL-22 and incubated overnight prior to RNA isolation. All retinoid-containing treatments were protected from light throughout the experiment.

### Liquid chromatography–ultraviolet spectroscopy and mass spectrometry measurement of retinol

Serum was collected from the abdominal aorta at the time of euthanasia and flash frozen. After thawing, serum samples were hydrolyzed in 50 mM potassium hydroxide at 37°C for 10 minutes.

HT-29 cells were cultured in 1X DMEM supplemented with GlutaMAX, 10% FBS, and 1% penicillin–streptomycin (ThermoFisher) at 37°C and 5% CO2. Cells were harvested after one to three days in culture by treatment with 0.25% trypsin-EDTA (Gibco), pelleted by centrifugation, and flash frozen.

Sample preparation for retinoid analysis was carried under yellow-light conditions. Samples were extracted twice with hexane under a yellow light, dried under nitrogen gas, and resuspended in methanol. For the LC-UV measurements, samples were analyzed using an Agilent 1290 Infinity II high-performance liquid chromatography system with an ultraviolet detector set to 325 nm. LC-MS measurements were performed using atmospheric pressure chemical ionization (APCI) on a SCIEX QTRAP 6500^+^ coupled to a Shimadzu LC-30AD HPLC system.

Fifteen microliters of sample were injected onto an Agilent InfinityLab Poroshell 120 EC-C18 column (2.1 × 150 mm, 2.7 μm) maintained at 35°C. The mobile phase consisted of solvent A (90% methanol, 10% water) and solvent B (100% methanol) at a flow rate of 0.7 mL/min. The elution gradient is shown in Table S2.

### Single cell RNA sequencing

Small intestines were recovered from vitamin A–replete and vitamin A–deficient mice at 8 weeks of age and flushed with sterile PBS. Intestines were opened longitudinally, cut into 2-cm segments, and washed three times in PBS containing 3% FBS and 0.04% BSA, followed by decanting of the supernatant. After the final wash, EDTA was added to a final concentration of 5 mM, and tissues were incubated at 37°C for 15 minutes. Cell suspensions were passed sequentially through 100-*μ*M and 70-*μ*M strainers on ice, pelleted by centrifugation at 300 *g* for 5 minutes, and resuspended in PBS containing 3% FBS and 0.04% BSA. Samples from each dietary condition were pooled prior to library preparation.

Library preparation and sequencing were performed in the UT Southwestern Microbiome Research Laboratory. Gel bead-in-emulsion (GEM) generation was carried out using a 10X Genomics Chromium Controller with the Chromium Next GEM Single Cell 3’ kit v3.1, Chromium Next GEM Chip G, and Dual Index Kit TT Set A. cDNA and barcoded sequencing libraries were prepared according to the manufacturer’s instructions and assessed for quality and concentration using a Bioanalyzer 2100 and qPCR, respectively.

Libraries passing quality control were sequenced using paired-end sequencing on a NextSeq 550 platform with a 150-cycle flow cell, generating approximately 25,000 to 30,000 reads per cell. Unique molecular identifier (UMI) counts were used to estimate the number of cells successfully captured and sequenced. Data were processed using the Cell Ranger software suite for demultiplexing, barcode processing, alignment to the mm10 mouse reference genome, and initial clustering.

Seurat was used for all downstream analyses of Cell Ranger-generated filtered matrices.^61^ Cells with fewer than 10 detected features, more than 2,000 detected features, or greater than 50% nuclear transcripts were excluded for quality control, as previously described.^62^ Data were normalized, scaled, and integrated with a pre-annotated adult mouse small intestine reference dataset from the Mouse Cell Atlas.^24^ Differential gene expression analysis was performed using the limma R package, and data were visualized using Seurat v5 and the ggplot2 package.

### Quantitative real-time PCR

For mouse tissue, terminal small intestine was collected and homogenized in MP Biomedical Lysing Matrix S tubes (Fisher) using an MP Biomedical FastPrep bead beater. Homogenates were centrifuged for 5 minutes prior to RNA extraction using the RNeasy Mini Kit (Qiagen) on a QIAcube instrument (Qiagen).

For cultured cells, monolayers were washed with PBS, lysed directly in Qiagen RLT buffer, and RNA was extracted using the same kit and platform. cDNA was synthesized from total RNA using an M-MLV–based reverse transcription protocol (ThermoFisher). cDNA was diluted 1:10 and used as template for TaqMan Expression Assays (see Key Resources Table) run on a QuantStudio 7 Flex Real-Time PCR System (Applied Biosystems). Relative gene expression was calculated using the comparative CT method, normalized to 18*S* rRNA.

### Immunoblotting

Small intestinal tissue was suspended in RIPA buffer supplemented with protease inhibitors and homogenized by rotor and stator. Homogenates were centrifuged, and supernatants were collected for analysis. Total protein concentration was determined by Bradford assay. Equal amounts of protein were resolved by SDS-PAGE and transferred to PVDF membranes. Membranes were blocked for 1 hour at room temperature in 5% blotting grade non-fat milk (Bio-Rad) in TBS-T and incubated overnight at 4°C with anti-REG3G antibody^4^ at a 1:1000 dilution in blocking buffer. Membranes were washed five times with TBS-T and incubated with goat anti-rabbit HRP-conjugated secondary antibody (Abcam, 1:5000) for 1 hour at room temperature. After five washes, membranes were imaged using a Bio-Rad ChemiDoc system. Membranes were then stripped (Thermo Scientific), then blocked and re-probed with anti-ACTB antibody (1:1000) as a loading control.

### Brightfield microscopy of mouse small intestine

Mouse small intestines were flushed with PBS and fixed by incubation in Bouin’s fixative (Ricca Chemical Company) with gentle rocking overnight at 4°C. Tissues were washed three times with 70% ethanol, embedded in paraffin, sectioned, and stained with hematoxylin and eosin at the UT Southwestern Histology Core. Slides were imaged using a Keyence BZ-X8000 microscope. Paneth cells were identified based on their position at the base of crypts and characteristic morphology and were enumerated manually.

### Immunofluorescence microscopy to visualize REG3G protein

Mouse small intestines were flushed with PBS and fixed in Bouin’s fixative overnight at 4°C with gentle rocking. Tissues were washed three times in 70% ethanol, embedded in paraffin, and sectioned at the UT Southwestern Histology Core. Sections were deparaffinized by incubation in xylene for 10 minutes twice, followed by rehydration through a graded ethanol series (100%, 100%, 95%, 95%, 90%, 90%, 70%) and final immersion in deionized water. Antigen retrieval was performed by boiling slides in citrate-based antigen retrieval buffer (Electron Microscopy Sciences) for 10 minutes, followed by cooling at room temperature for 20 minutes.

Sections were blocked with CAS blocking buffer (ThermoFisher) for 1 hour at room temperature and incubated overnight at 4°C with anti-REG3G antibody (1:50).^4^ Rabbit polyclonal IgG was used as an isotype control. Slides were washed three times with PBS and incubated with goat anti-rabbit Alexa Fluor 594-conjugated secondary antibody (1:500; Abcam) for 1 hour at room temperature, protected from light. Slides were washed three times with PBS and mounted using Fluoromount-G Mounting Media with DAPI (SouthernBiotech 0100-20) to label nuclei. Images were acquired using a Keyence BZ-X800 microscope, and fluorescence intensity was quantified using Fiji software.

### Immunofluorescence microscopy to visualize spatial separation of luminal microbes and intestinal villi

Mouse small intestines were dissected, and luminal contents were undisturbed. Tissues were fixed in a 60% methanol, 30% chloroform, 10% glacial acetic acid solution, and then washed with 70% ethanol for at least 24 hours before paraffin embedding and sectioning at the UT Southwestern Histology Core. Slides were deparaffinized with two 10–minute incubations of xylene followed by a rehydration ethanol gradient (100%, 100%, 95%, 95%, 90%, 90%, 70%) for 5 minutes each. Finally, slides were placed in distilled water and then blocked with 5% bovine serum albumin (BSA) in PBS-T (0.1% Tween 20 diluted in 1X PBS) for one hour at room temperature. Slides were then incubated with a 1:100 dilution of anti-UEA-1 (EY Labs) and anti-Lipid A (GeneTex) in 2% BSA PBS-T overnight at 4°C. Slides were washed three times with PBS-T and then incubated with streptavidin conjugated with Alexa Fluor 488 (1:500) and donkey anti-goat Alexa Flour 594 (1:500) for one hour at room temperature, protected from light. Slides were washed three times with PBS-T and mounted using DAPI Fluoromount (SouthernBiotech 0100-20) and imaged on a Keyence BZ-X800 microscope.

### Chromatin immunoprecipitation (ChIP)

To isolate small intestinal epithelial cells, mouse small intestines were flushed with PBS, opened longitudinally, and washed twice more in PBS. Tissue was cut into 1-cm pieces and washed by brief vortexing in PBS. Samples were incubated in 5 mM EDTA in PBS at 37°C for 10 minutes with shaking at 150 rpm, followed by an additional 10-minute incubation at 4°C with gentle rocking. Tissue suspensions were briefly vortexed and passed sequentially through 100-*μ*m and 70-*μ*m strainers. Cells were pelleted by centrifugation at 650 *g* for 5 minutes at 4°C and resuspended in PBS. ChIP CrossLink Gold (Diagenode) was added directly to a final concentration of 0.4% and incubated for 45 minutes. Cells were washed with PBS and then fixed in 4% paraformaldehyde for 15 minutes at room temperature with rotation. Crosslinking was quenched by adding glycine to a 100 mM final concentration. Fixed cells were snap frozen and stored at -80°C until further processing.

For ChIP analysis of cultured cells, HCT-116 cells were grown in 175-cm^2^ flasks to ∼80% confluence then treated overnight with 1 *μ*M retinol and 100 ng/uL of recombinant IL-22 (Abcam). Media was aspirated, and cells were washed twice with PBS. Cells were incubated with 0.4% ChIP CrossLink Gold in PBS for 30 minutes, washed twice with PBS, and fixed in 4% paraformaldehyde for 15 minutes at room temperature with gentle rocking. Crosslinking was quenched by adding glycine to a 100 mM final concentration.

Immunoprecipitation of fixed mouse or human cell pellets was performed using the iDeal ChIP-seq kit for Transcription Factors (Diagenode), following the manufacturer’s protocol. Chromatin was sonicated using a Bioruptor Pico sonicator (Diagenode) for six cycles of 30 seconds on and 30 seconds off. Nuclear lysates were immunoprecipitated with equal volumes of anti-RARα antibody, anti-STAT3 antibody, or rabbit IgG as an isotype control. Recovered DNA was quantified by qPCR using region-specific primers (see Key Resources Table) and SYBR Green reagents (ThermoFisher) and measured on a QuantStudio 7 Flex Real-Time PCR System (Applied Biosystems).

### Luciferase reporter assays

A 1313-base pair fragment of the human *REG3G* promoter (-976 to +337 relative to the transcription start site) was cloned upstream of the firefly luciferase gene in a pGL2 plasmid backbone using KpnI and XhoI restriction sites. A synthetic RAR-responsive reporter plasmid (Addgene) and pCMV-LacZ (Takara).^63^ Site-directed mutagenesis was performed using region-specific primers (see Key Resources Table). The full-length sequence of either *GFP* or *RAR403* was cloned into the overexpression pcDNA3.1+ plasmid backbone using EcoRI and XhoI restriction enzymes.

HEK293 cells were transfected using OptiMEM Reduced Serum Media (Gibco) and Fugene HD transfection reagent (Promega). Cells were incubated for 24 hours prior to lysis in Passive Lysis Buffer (Promega). Luminescence was measured using Bright-Glo reagents (Promega) on a BioTek Synergy LX multimode reader (Agilent). β-galactosidase activity from the pCMV-LacZ transfection control plasmid was measured using a β-Galactosidase Enzyme Assay System (Promega) and quantified on the same instrument to normalize for transfection efficiency.

### siRNA knockdown

HCT-116 cells were cultured in DMEM (1X) supplemented with GlutaMAX, 10% fetal bovine serum, and 1% penicillin-streptomycin (ThermoFisher) at 37°C with 5% CO2. At approximately 80% confluence, cells were transfected with pooled siRNAs targeting *RARA*, *RARG*, or with a nontargeting control siRNA (see Key Resources Table), using Lipofectamine RNAiMax (ThermoFisher) as a transfection agent. Twenty-four hours after transfection, cells were provided with fresh medium containing 1 *μ*M retinol (Sigma-Aldrich) and/or 100 ng/mL of IL-22 (Abcam). After an additional 24 hours, cells were washed with PBS and RNA was extracted using the Qiagen RNeasy Mini Kit (Qiagen) on a QIAcube instrument.

### Computational Motif Mapping

To identify candidate transcription factor binding sites within the REG3G promoter, the nucleotide sequence of the promoter region was analyzed using the Bioconductor R package TFBSTools.^56^ JASPAR position weight matrices corresponding to human and mouse RARα and STAT3 were retrieved from the JASPAR 2024 database and used to scan the promoter sequence. Putative binding sites were scored based on relative match strength to each matrix, allowing identification of the highest confidence predicted transcription factor binding motifs.

### Statistical analysis

Detailed information regarding statistical analyses, including definitions of significance and the specific statistical tests used, is provided in the figure legends. Data are presented as mean ± standard error of the mean (SEM). Sample size was not determined by formal power calculations. Investigators were not blinded to group allocation. No formal randomization strategy was employed; however, animals were assigned to experimental groups in a random manner, and samples were processed without a predetermined order. Statistical analyses were carried out using GraphPad Prism and, when appropriate, R software.

## Key Resources Table

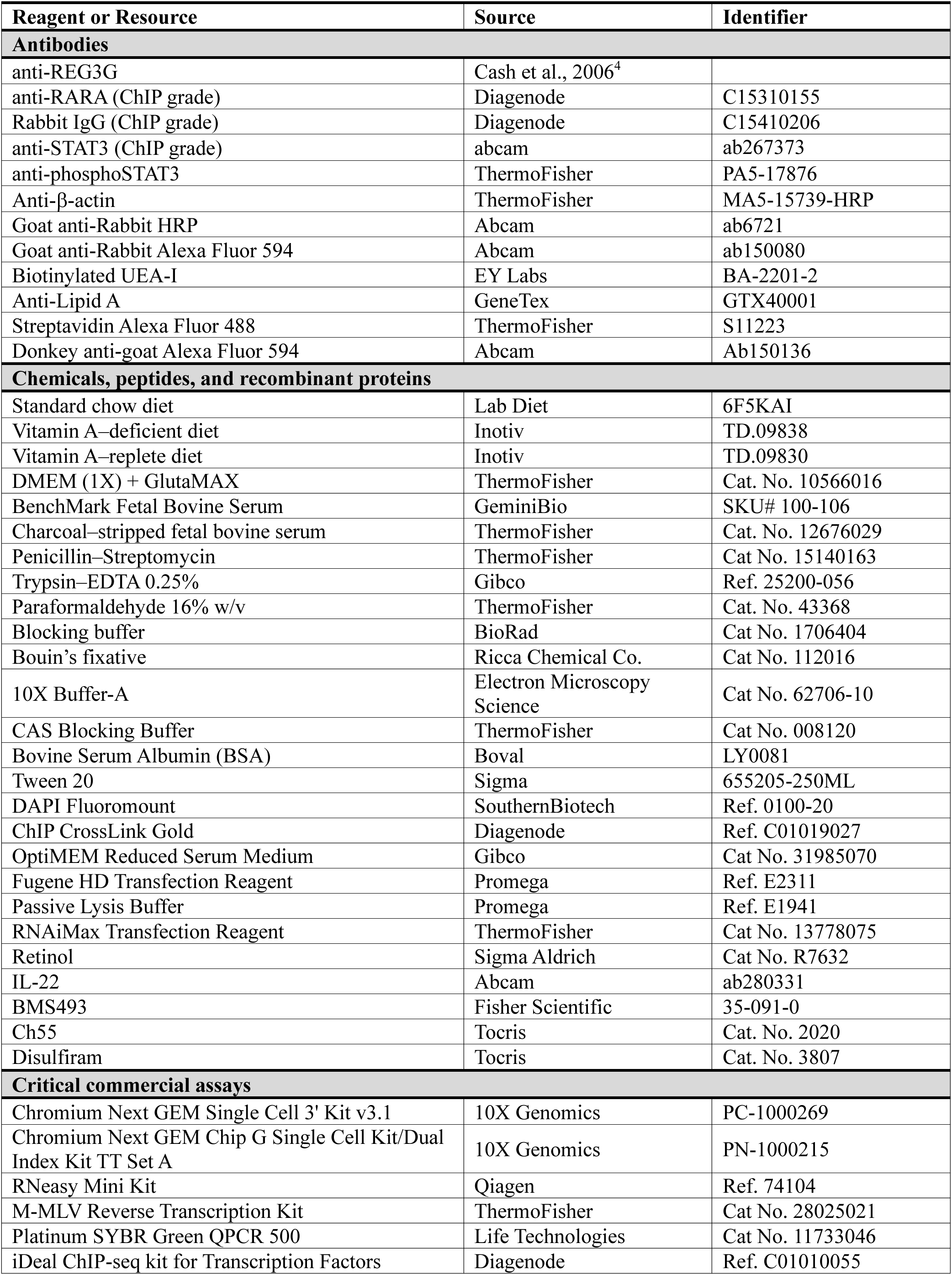
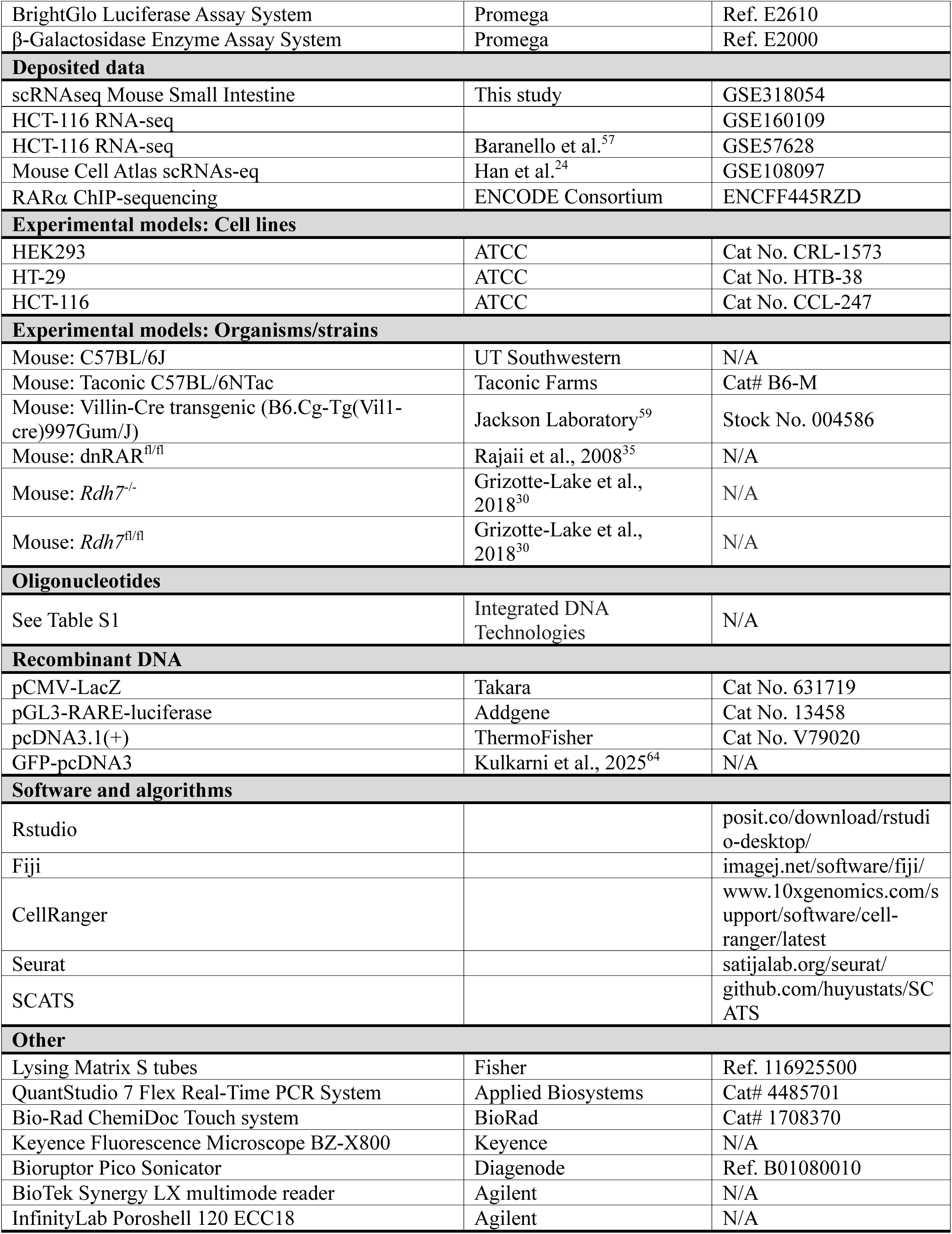

## Supplementary tables

**Table S1:**
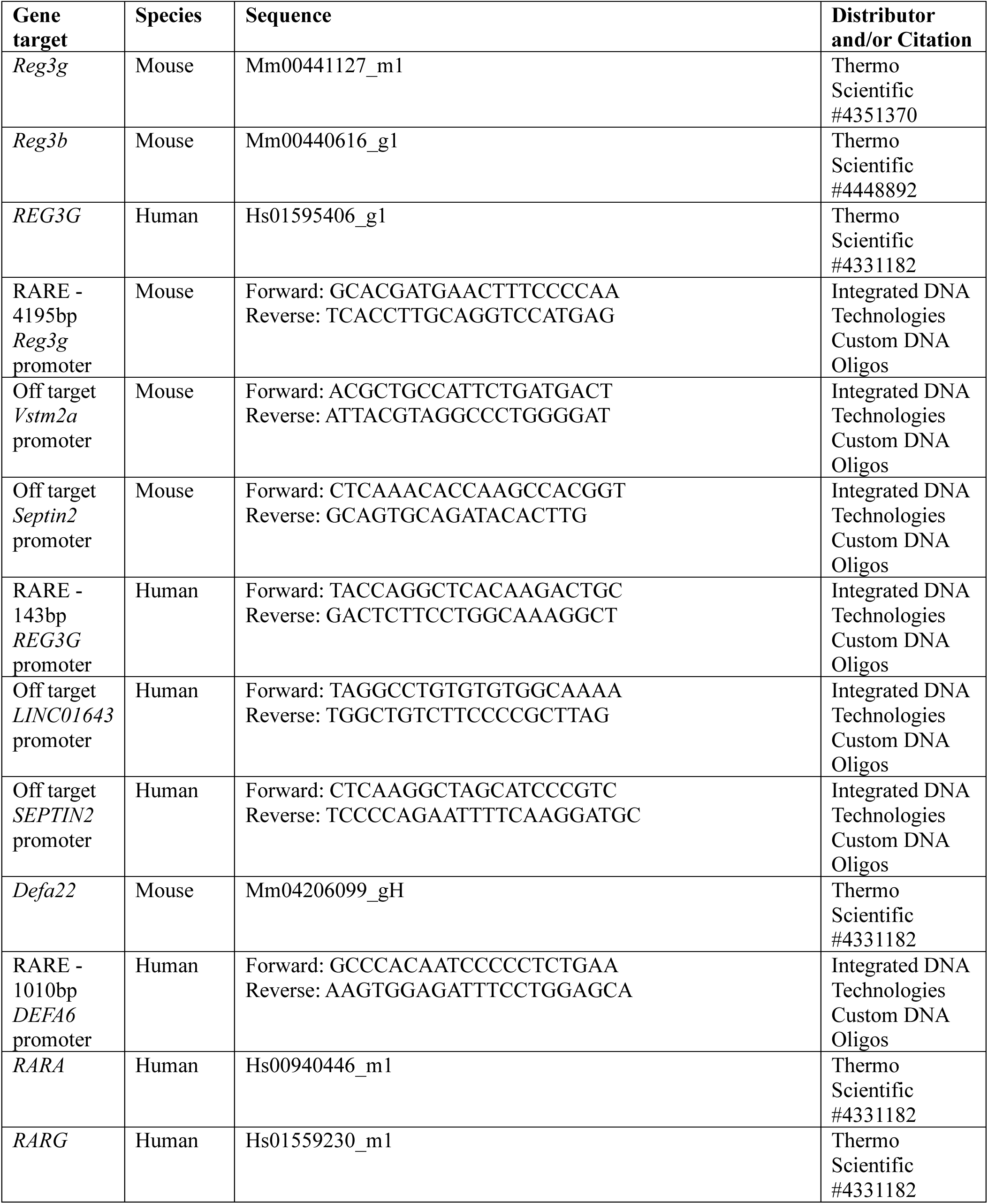
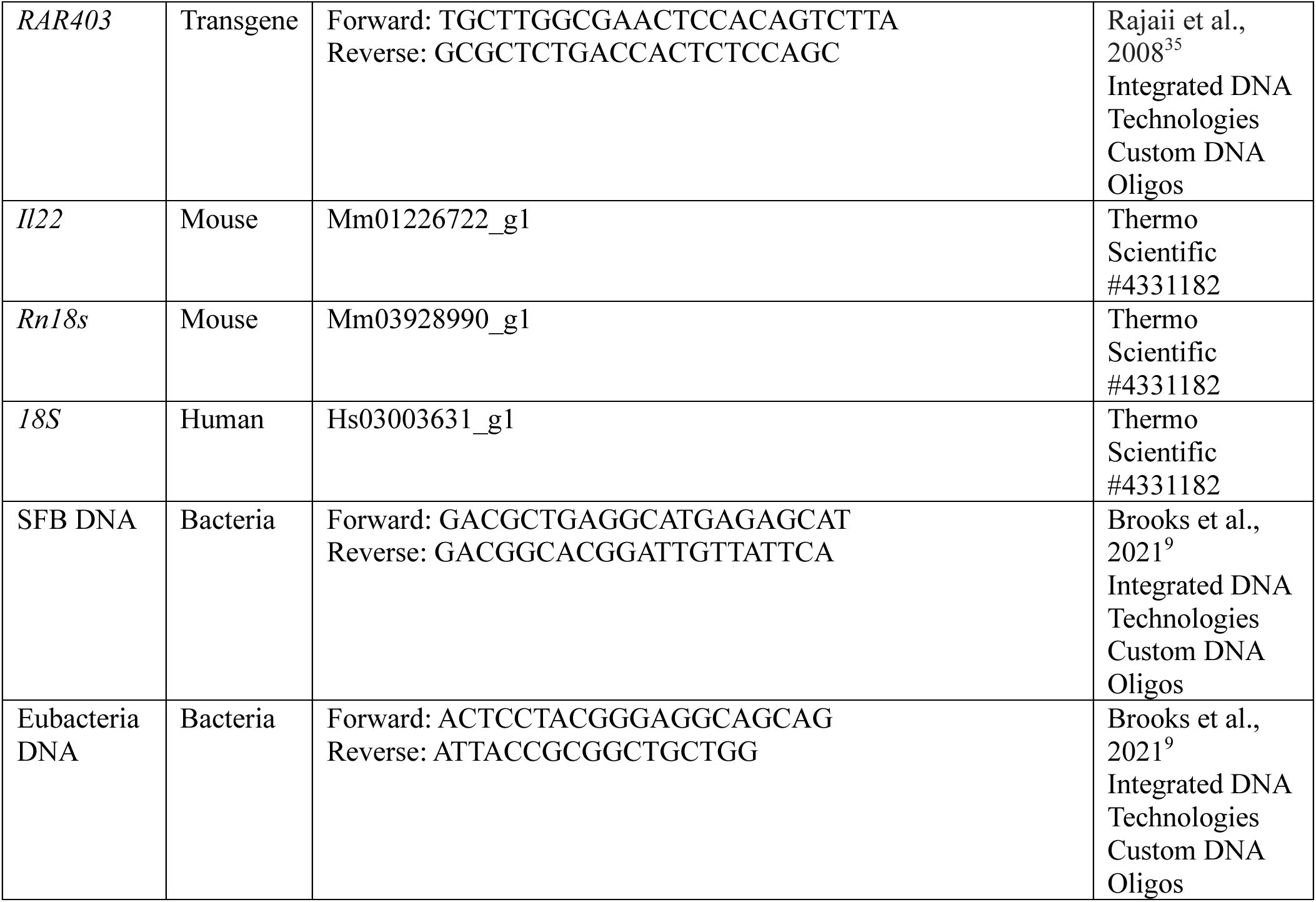
Oligonucleotides and primers.

**Table S2:** LC-UV gradient program for retinol detection.

| Time (min) | Mobile A (%) | Mobile B (%) |
| --- | --- | --- |
| 0.0 | 100 | 0 |
| 2.5 | 0 | 100 |
| 8.5 | 0 | 100 |
| 8.6 | 100 | 0 |
| 10.0 | 100 | 0 |

